# TDP-43 dysfunction leads to impaired proteostasis and predisposes mice to worse neurological outcomes after brain injury

**DOI:** 10.1101/2025.10.20.683438

**Authors:** Melissa S. Rotunno, Megan Fowler-Magaw, Jianjun Zhong, Kennedy O’Hara, Elenore A. Wiggin, Debra Cameron, Karly Stallworth, James Bouley, Holly McEachern, Mina N. Anadolu, Jeffrey A. Nickerson, Andrew R. Tapper, Susanna Molas, Francesca Massi, Nils Henninger, Oliver D. King, Daryl A. Bosco

## Abstract

**Background:** Pathological TAR DNA-binding protein 43 (TDP-43) dysfunction is associated with multiple neurodegenerative disorders. However, the mechanistic link between TDP-43 dysfunction and neurodegeneration is poorly understood and likely involves a combination of genetic and environmental risk factors. A major risk factor for neurodegenerative disease is exposure to traumatic brain injury (TBI). Here, we investigated the synergistic interplay between TDP-43 dysfunction and TBI in a murine model of amyotrophic lateral sclerosis (ALS)/frontotemporal dementia (FTD).

**Methods:** A model of TDP-43 dysfunction caused by a knock-in Q331K mutation in *Tardbp* was combined with a mild model of TBI. Control conditions included both WT mice and mice with sham surgery. Animals were evaluated for behavioral deficits at timepoints pre- and post-surgery. Additionally, post-mortem brain tissues were examined using RNA sequencing and mass spectrometry-based quantitative proteomics together with histological and biochemical analyses.

**Results:** Expression of dysfunctional TDP-43 *in vivo* caused deficits in multiple branches of the proteostasis network, including protein folding, protein synthesis, and protein turnover. Examples include mis-expression of chaperones and genes within the ubiquitin-proteosome pathway in mutant TDP-43 versus WT mice. Further, mutant TDP-43 expression correlated with reduced thermostability of proteins associated with the ribosome and the chaperonin containing TCP-1 complex. In response to TBI, mutant TDP-43 mice exhibited significantly worse neurological outcomes relative to WT animals. Heightened neurological deficits in mutant TDP-43 mice following TBI coincided with a robust upregulation of proteostasis- and stress-related genes at the transcript level. However, this upregulation was not detected at the protein level.

**Conclusions:** Our data demonstrate that expression of dysfunctional TDP-43 leads to deficits within the proteostasis network in vivo at baseline. Despite an upregulation of proteostasis-related genes at the transcript level in mutant TDP-43 mice after TBI, mutant TDP-43 mice exhibit an impaired response to, and recovery from, brain trauma relative to their WT counterparts. Restoring proteostasis is expected to protect against the detrimental effects of TDP-43 dysfunction, especially under stress conditions that promote neurodegenerative disease.

## BACKGROUND

TAR DNA-binding protein 43 (TDP-43) is an essential RNA/DNA-binding protein that performs critical functions in the context of RNA splicing, transport, and expression.(1) TDP-43 shuttles between the nucleus and cytoplasm while fulfilling these functions, although the protein is predominately expressed in the nucleus under physiological conditions. A significant cytoplasmic shift in TDP-43 localization has been observed in cultured cells exposed to various external stimuli or “stressors” that inhibit proteasomal and transcriptional activity, or that induce hyperosmolarity, hyperexcitability or excitotoxicity.(2–6) While the mechanisms underlying stress-induced cytoplasmic accumulation of TDP-43 have not been fully elucidated, recent reports have shown that both RNA binding capacity and post-translational modification of TDP-43 affect TDP-43 subcellular localization.(3, 7–10) TDP-43 has also been identified as a component of cytoplasmic condensates called stress granules, which form in response to acute cellular stress.(11–13) Autosomal dominant mutations in TDP-43 that cause familial amyotrophic lateral sclerosis (ALS) alter the properties of stress granules as well as other TDP-43-associated ribonucleoprotein (RNP) granules,(10–12, 14–16) possibly through altered phase separation behavior of mutant TDP-43 with itself, other RNA-binding proteins and/or RNA.(10, 17–20) Collectively, these observations indicate that subcellular TDP-43 localization can serve as a proxy for cellular stress. Further, the effects of TDP-43 on stress response pathways can be perturbed by disease-causing mutations.

In addition to TDP-43 mutations in familial ALS, TDP-43 has emerged as a hallmark pathological feature across multiple neurodegenerative disorders termed TDP-43 proteinopathies,(21) including ALS, frontotemporal lobar degeneration (FTLD),(22, 23) Alzheimer’s disease (AD),(24) limbic-predominant age-related TDP-43 encephalopathy (LATE)(25) and chronic traumatic encephalopathy (CTE).(26). Cells with cytoplasmic accumulation and/or nuclear depletion of TDP-43 have been shown to exhibit splicing defects that are consistent with TDP-43 loss-of-function in human post-mortem CNS tissues from ALS/FTLD and AD cases.(4, 27–31) Most cases of human TDP-43 proteinopathy occur sporadically with poorly defined etiologies, and likely develop through a combination of genetic susceptibility and environmental risk factors.(32, 33) Indeed, emerging evidence supports that ALS pathogenesis occurs through a multistep process, even in cases with penetrant genetic mutations in *TARDBP* encoding TDP-43.(34) The multistep hypothesis purports that individual ALS-linked mutations cannot fully account for the disease process, which must include other factors such as additional genetic mutations, environmental exposures and/or physical stress.(34, 35)

Traumatic brain injury (TBI) is a form of physical stress that also represents a major risk factor for developing neurodegenerative diseases, including those with TDP-43 pathology.(36) Cytoplasmic shifts in TDP-43 localization have been reported in laboratory animal models of TBI.(13, 37–41) Further, recurrent TBI leads to CTE with cytoplasmic TDP-43 pathology in humans.(26, 42) Here, we sought to investigate the relationship between stress and TDP-43 dysfunction in the context of a multistep model of neurodegenerative disease. To this end, we combined a murine model of TDP-43 dysfunction caused by a genetic knock-in mutation in *Tardbp* together with a mild model of TBI.(43, 44) Our results revealed that mutant TDP-43 mice exhibit significantly worse neurological outcomes during the acute phase post-TBI relative to their wild-type (WT) counterparts. Corresponding transcriptomics and RT-PCR analyses revealed synergistic effects of TDP-43 mutation and TBI on gene expression and processing. For example, a significant upregulation of proteostasis- and stress-related genes was detected specifically in brain tissue from mutant TDP-43 mice after TBI. Many of the pathways that became upregulated in mutant TDP-43 mice following TBI were also highlighted in a recent transcriptomics analysis of post-mortem ALS and FTLD-TDP CNS tissues.(45, 46) While this TBI model caused a cytoplasmic shift in TDP-43 localization, this change in TDP-43 localization did not appear to correlate with mis-splicing events caused by TDP-43 loss-of-function. Rather, we found that mutant TDP-43 expression correlated with reduced thermostability of cytoplasmic ribosomal proteins. In fact, diminished thermostability of ribosomal proteins correlated with mutant TDP-43 expression, irrespective of whether mutant TDP-43 mice had been exposed to brain trauma or not. Notably, TDP-43 has been shown to associate with ribosomes and to modulate both global and local translation.(47–50)

The outcomes of this study are consistent with a model in which expression of mutant TDP-43 leads to deficits throughout the proteostasis network. Upon CNS challenge, compensatory mechanisms appear to become activated in mutant TDP-43 mice, yet these mice fail to mount the physiological response to brain injury exhibited by their WT counterparts. Therefore, our data demonstrate that expression of dysfunctional TDP-43 leads to impaired response to, and recovery from, brain trauma and provide mechanistic insight into TBI as a risk factor for TDP-43 proteinopathies. Restoring proteostasis may therefore protect against the effects of TDP-43 dysfunction, especially under conditions of CNS challenge that promote neurodegenerative disease.

## METHODS

### Animals

All animal related protocols were conducted in accordance with the Guide for the Care and Use of Laboratory Animals published by the National Research Council (US) Committee. The protocols were approved by the Institutional Animal Care and Use Committee (IACUC) of the University of Massachusetts Chan Medical School. Mice with the human-equivalent TDP-43 Q331K mutation on a pure C57BL/6J (WT) background were generated and characterized in a previous study.(44) All experiments herein were performed with aged-matched, male mice with either the homozygous TDP-43 Q331K or TDP-43 WT genotype. Mice were generated using standard breeding approaches or via *in vitro* fertilization at the UMass Chan Medical School Transgenic Animal Modeling Core (TAMC; Worcester, MA). All mice were maintained under standard conditions with no more than 5 mice per cage, on a standard 12h light/dark cycle (white light ON at 7:00am and OFF at 7pm), and were provided *ad libitum* access to food and water.

### Traumatic brain injury (TBI) surgery

Mice at 16-weeks of age were exposed to TBI or sham surgery. TBI surgeries were based on a previously published protocol.(43) Details of the TBI surgery used herein were published by us elsewhere.(51) Briefly, mice were anesthetized with 3% isoflurane (Pivetal, NDC 46066-755-03) in carrier gas at 1.5 L/min flow. Prior to surgery, buprenorphine (PAR Pharmaceutical, NDC 42023-179-05) was administered subcutaneously at 1 mg/kg. Hair was removed from the top of the head and the scalp skin was disinfected with 75% ethanol and betadine. A ∼1.2-1.5 cm incision was made along the midline starting approximately 3 mm posterior from the eyes to expose the right parietal skull. The skin over the left and right hemispheres of the skull was excised. The periosteum was removed and the exposed skull was positioned on a cushion beneath the weight-drop. A transducer rod tip was aligned to the skull-top location: 2.5 mm posterior to Bregma and 2 mm lateral to the midline. From a height of 15 cm, a 50-gram weight was released that came into contact with a transducer, which in turn delivered an impact onto the mouse’s intact skull (see diagram in **Fig. 1A**). After the impact, mice were placed onto a heating blanket in a supine position for the time needed to wake from the anesthesia (i.e., time required to right themselves to a prone position), which took on the order of minutes for each mouse. Mice were then returned to the anesthesia chamber in preparation for a subsequent impact; this process was repeated a total of ten times within a single surgical session. As the impacts were delivered within the same surgical session and not spaced by days or weeks, this model is not considered a repetitive TBI model that we described previously.(40, 52) After the surgery was completed, cefazolin (HIKMA Pharmaceutical, NDC 0143-9924-90) was administered intramuscularly at 500 mg/kg and meloxicam (Norbrook, NDC 55529-040-10) was administered subcutaneously at 5 mg/kg. Mice were placed on a heating blanket and monitored until they were ambulatory (approximately 15 minutes), and then transferred to their home cage. Buprenorphine was administered as described above at 8 h, 16 h and 24 h after the first dose. For the sham surgery, mice underwent the same procedure as the TBI mice but without the weight-drop impact.

**Figure 1.**
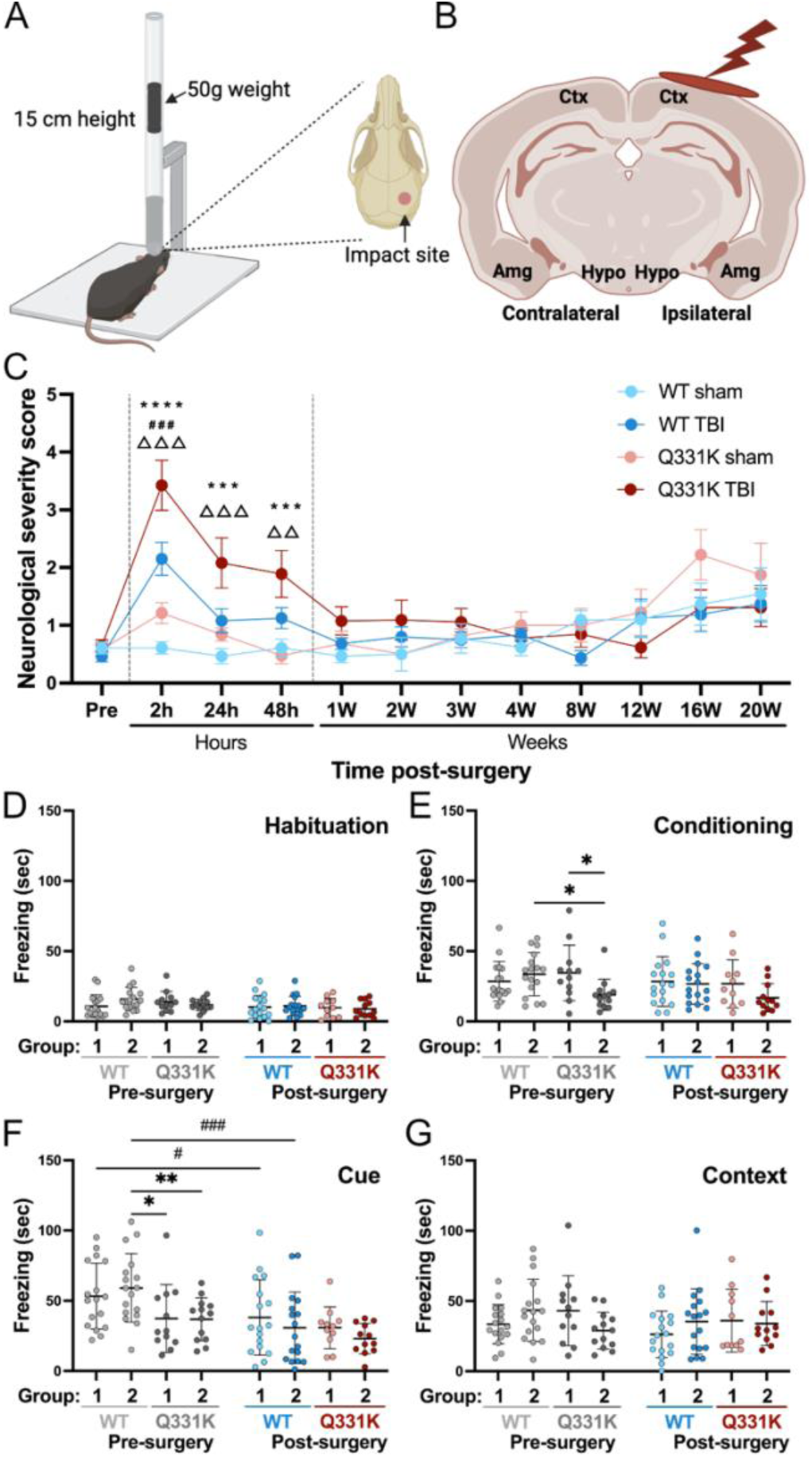
Neurological and cognitive deficits in mice with and without brain trauma. **(A)** Diagram of the weight drop device used to induce multiple closed-skull impacts with a 50g weight dropped from a 15cm height within a single surgical session. Location of impact site (red circle) is shown on the right. Created with biorender.com. **(B)** Illustration depicting the location of the impact site (red lightning bolt and oval). Several brain regions of interest are highlighted as follows: cortex (Ctx), amygdala (Amg), and hypothalamus (Hypo) on both ipsilateral and contralateral sides of the brain. Created with biorender.com. **(C)** Neurological severity score (NSS) plotted as a function of time post-surgery and cohort (see key). Statistically significant comparisons shown on the graph include Q331K TBI versus Q331K sham at 2h (****p=2.98e^-6^), 24h (***p=0.00058) and 48h (***p=0.00018) post-surgery, WT TBI versus WT sham at 2h post-surgery (^###^p=0.00070) and Q331K TBI versus WT TBI at 2h (^ΔΔΔ^p=0.00026), 24h (^ΔΔΔ^p=0.00022) and 48h (^ΔΔ^p=0.0058) post-surgery. Additional significant comparisons include Q331K TBI versus WT sham at 2h (p=7.0e^-10^), 24h (p=2.54e^-6^) and 48h (p=0.00014) post-surgery. **(D-G)** Mice were randomly assigned to Group 1 (sham) or 2 (TBI). Results of the cued and contextual fear conditioning (CCFC) test. **(E)** The Q331K-2 cohort exhibited altered freezing behavior during conditioning compared to Q331K-1 (*p=0.028) and WT-2 (*p=0.017). **(F)** Both Q331K cohorts exhibited attenuated cue-induced freezing relative to WT animals (*p=0.012 for WT-2 vs. Q331K-1; **p=0.009 for WT-2 vs. Q331K-2) pre-surgery. WT animals exhibited attenuated cue-induced 5-months after sham (^#^p=0.048) and TBI (^###^p<0.001) surgeries.

### Neurological Severity Score (NSS) Test

The NSS test was performed at 12 time points: pre-surgery and 2h, 24h, 48h, 1 week, 2 weeks, 3 weeks, 4 weeks, 8 weeks, 12 weeks, 16 weeks and 20 weeks post-surgery, as previously described in detail.(43, 53) Body weight was measured prior to every NSS testing session. Briefly, the NSS consists of 10 individual tasks designed to assess motor function, alertness and physiological behavior. Each task was completed regardless of whether the mouse passed or failed previous tasks. For each task, mice were assigned a score of 1 to indicate failure of the task or 0 if the task was completed, resulting scores ranging between 0 (no deficit) and 10 (maximum deficit). The NSS test was performed by a lab technician who was blinded to the genotype and surgery type of the mice. For statistical analyses, between-group comparisons of the NSS over repeated measurements (time) were conducted using longitudinal mixed models. Time was treated as a categorical variable. The models included group (each group representing both a genotype and surgery type) and time as fixed covariates, as well as the two-way interaction. When the main effects were significant, pairwise comparisons were conducted to identify significance by group and time point. A two-sided p<0.05 was considered statistically significant. Statistical analyses were performed using SPSS Statistics Version 29 (IBM-Armonk, NY).

### Cued and Contextual Fear Conditioning (CCFC) Test

Animals were generated through two in vitro fertilization procedures at the Transgenic Animal Modeling Core (TAMC) at UMass Chan Medical School using heterozygous Q331K male and female mice. Homozygous Q331K and WT mice were randomly assigned to group 1 or group 2, and both groups were subjected to the CCFC test before surgery. The CCFC test was conducted at the UMass Chan Mouse Behavioral Core facility using the EthoVision XT system (Noldus, BOM-EV-0001). Mice were placed in a Ugo Basile Fear Conditioning Chamber (Noldus, BOM-XUGO-F101) made of optical transparent plastic 26 cm long, 26 cm wide and 38 cm tall. The top of the chamber was equipped with a monochrome GigE camera (Noldus, BOM-XCAM-GIGE) used to track animal movement in real-time during the testing session. The bottom of the chamber contained electricity-conductive metal strips to deliver foot shocks. The chamber was nested in a cabinet equipped with a speaker to deliver white noise and high-frequency tones per the experimental design detailed below. The cabinet wall was insulated to reduce environmental noise and a built-in ventilator allowed for air-exchange. All the inputs, including white noise, tone, and foot shock, were controlled with a computer-based operation console (Ugo Basile, version 1.1.8.9). EthoVision XT 15 software was used for animal tracking and data analysis. The live tracking started immediately when the animal entered the chamber, after which there was a 60s delay prior to the start of an input. The full test session was conducted over four consecutive days as follows. Note that the white noise and the tone input were delivered randomly, with random intervals (ranging from 5s to 30s) between these two inputs for all tests on each day.

Day 1:

White noise: 50% intensity, 30s duration, and repeated 5 times.

Tone: 1000 Hz, 50% intensity, 30s duration, and repeated 5 times.

Day 2:

White noise: 50% intensity, 30s duration, and repeated 5 times.

Tone: 1000 Hz, 50% intensity, 30s duration, and repeated 5 times.

Electrical foot shock: 0.75 mA intensity that occurred at the last second of each tone. Each mouse received five foot shocks in total on Day 2 with each shock lasting for 1s.

Day 3:

White noise: 50% intensity, 30s duration, and repeated 5 times.

Tone: 1000 Hz, 50% intensity, 30s duration, and repeated 5 times.

Novel context: To create a novel context, a new scent and visual component were introduced to the chamber as follows. One drop of vanilla extract (novel scent) was applied to a piece of tissue paper, which was placed in the feces tray underneath the chamber. Paper boards with white and black stripes (novel visual) were attached to the outer side of the chamber.

Day 4:

The context was reverted as per Days 1 and 2 (i.e., the paper boards were removed and the feces tray cleaned before Day 4) for testing on Day 4.

Neither white noise nor tone were implemented.

Below is a table summarizing the inputs and contexts for the CCFC test:

|  | Day 1 | Day 2 | Day 3 | Day 4 |
| --- | --- | --- | --- | --- |
| White noise | + | + | + | - |
| Tone | + | + | + | - |
| Foot shock | - | + | - | - |
| Novel context | - | - | + | - |

For analysis of Days 1-3, the initial 29s of each input was extracted to avoid potential interference from the last second when the foot shock was delivered. Therefore, the total duration for white noise and tone inputs is 29s x 5 = 145s. For Day 4, the consecutive 145s after the initial 60s delay was extracted for the analysis. Mouse movement was detected by the software at the pixel level for the entire chamber from one video frame to the next. Thresholds based on animal activity within the chamber were set as follows: immobility state (<0.1% pixel change in the chamber), movement between immobility and walking (between 0.1% and 0.5%) and walking (>0.5%). The amount of time animals spent in an immobility state was interpreted as freezing behavior and plotted for each day (Day 1-4). The statistical analysis was performed as described above for NSS, where between-group comparisons over repeated measurements (time) were conducted using longitudinal mixed models. Time was treated as a categorical variable. The models included group and time as fixed covariates, as well as the two-way interaction. When the main effects were significant, pairwise comparisons were conducted to identify significance by group and time point. A two-sided p<0.05 was considered statistically significant. Statistical analyses were performed using SPSS Statistics Version 29 (IBM-Armonk, NY).

### RNA isolation

Mouse brain tissue samples (∼40 mg and 4 mm of diameter) were collected from the impact site (**Fig. 1A**) and the corresponding site on the contralateral side using a sterile disposable biopsy punch (Integra Miltex, 33-32) that excised tissue from the brain surface (i.e., cortex) to the brain bottom (i.e., hypothalamus). Total RNA was isolated from mouse brain tissues using Qiagen’s RNeasy Lipid Tissue Mini Kit (Cat. No. 74804) with DNase treatment following the manufacturer’s protocols. RNA concentration and absorbance were measured by a Nanodrop device (ND1000). Samples with A260/A280 between 1.8 to 2.2 were considered suitable for downstream applications.

### Transcriptome analysis by RNA-sequencing (RNAseq)

For the 24h-post TBI timepoint, RNAseq analysis was performed on RNA isolated from brain tissue as described above for six groups of mice: homozygous Q331K and WT genotypes, each under naïve, sham, and TBI conditions. For the 7mo-post TBI timepoint, only sham and TBI conditions for homozygous Q331K and WT genotypes were included. Sham and TBI conditions are described under TBI surgery. Naïve mice were not subjected to surgery and are therefore treated a separate control group than sham surgery mice. Between 3-6 mice were included per group for a total of 46 samples. RNA quality assessment, library preparation and sequencing were conducted by Novogene Co., LTD (Beijing, China). Poly-T oligo-attached magnetic beads were used for poly-A capture, random hexamer primers for first strand cDNA synthesis, and dTTP for second strand cDNA synthesis of non-directional libraries. Libraries were sequenced to a depth of 50M-125M raw read pairs (2x150 PE) per sample using an Illumina NovaSeq 6000 sequencer. After trimming adapter sequences and low-quality bases, reads were mapped with HISAT2 (2.0.5)(54) to the mouse mm39 reference genome with Ensembl release 107 gene annotations. Two samples (one Q331K sham and one Q331K TBI) from the 24h post-TBI timepoint were excluded from further analyses, as they were strong outliers in Principal Component Analysis (PCA) of expression values (counts-per-million on log_2_ scale with prior.count=1; **Fig. S1A**); see **Additional File 1** for all supplemental figures. These samples were also outliers for several mapping quality control (QC) metrics, with higher-than-usual fractions of multi-mapping, exonic, and unspliced reads. Two samples (one Q331K sham and one WT TBI) were also excluded from the 7mo-post TBI dataset because they were determined to by outliers in the PCA analysis (**Fig. S1B**).

Differential gene expression analyses were performed in R (4.3.2) using quasi-likelihood methods in the edgeR package (4.0.12).(55) Data from 24hr-post TBI and 7mo-post TBI experiments were analyzed separately. The TMM method was used for computing normalized library sizes and genes with low counts were filtered out in preprocessing. For both datasets, ∼25,000 genes were retained with at least 5 scaled counts in at least 3 samples, where the scaling adjusts for differences in total library sizes (read counts across all genes) between samples while keeping the median library size unchanged. The function glmQLFit (with non-default options robust=T and prior.count=1) was used to fit the expression data with a design matrix that included factors for injury (INJ, with levels Naïve, Sham, and TBI), genotype (GT, with levels Q331K and WT) and their interaction (model formula ∼0 + GT:INJ). Differential expression was assessed using the function glmQLFTest for contrasts that included nine simple between-group comparisons (WT.TBI vs WT.Sham, etc.); main effects of GT (Q331K vs WT) and INJ (TBI vs Sham), though in both cases excluding the groups with INJ=Naïve; and a test for an interaction between GT and INJ, again excluding the groups with INJ=Naïve. For the 7mo-post TBI data the naïve mice were not included, and the model also included an additive factor for IVF batch (with two levels, 162 and 167). Supplemental Table I includes the log_2_(fold-change) [LFC], p-value, and FDR for each gene for each contrast discussed in the results section.

Gene Set Enrichment Analysis (GSEA, 4.3.3) was performed using the Hallmark (mh.all.v2024.1.Mm.symbols.gmt) and Gene Ontology Biological Process (GOBP, m5.go.bp.v2024.1.Mm.symbols.gmt) gene set collections from the Molecular Signatures Database (MSigDB).(56–60) Cutoffs on Normalized Enrichment Score (NES) and adjusted p-value (FDR) were used to determine significantly altered gene sets (|NES| > 1.5 and FDR<0.05). NES is a score that reflects the overrepresentation of a gene set at the upper (positive value) or lower (negative value) end of a ranked gene list, after normalization to account for variation in gene set size. The gene list was pre-ranked based on directional p-values for the indicated comparison (-log_10_(p-value)*sign of log_2_(fold change)). A weighted enrichment statistic was used in the analysis. The 10 top gene sets based on highest |NES| are displayed for all enrichment analyses. If fewer than 10 are displayed then all gene sets that meet the cutoff criteria are shown. All gene set hits that meet the cutoff criteria can be found in **Supplemental Table II**.

### Quantitative Proteomics

Mice were perfused with PBS prior to tissue isolation for all proteomic analyses. Mouse brain tissue samples were collected as described above under RNA isolation and cut into ∼1 mm diameter pieces. For whole brain proteomics, the tissue was transferred to a pre-cooled glass homogenizer on ice with 900 μl of RIPA buffer (Sigma-Aldrich R0278-50ML) containing 1x protease inhibitor (Roche, 11873580001) and 1x phosphatase inhibitor (Roche, 04906837001). Complete tissue homogenization was performed with a WHEATON Power Homogenizer and Overhead Stirrer (catalog number 903475, serial number G082210) by setting the stirring speed on level 2 for 1 minute. The sample was transferred to a plastic tube together with 300ul of RIPA buffer that was added to rinse the homogenizer. The sample was left on ice for 30 min and then centrifuged at 15,899 xg for 10 min at 4°C in a table-top centrifuge (AXYGEN, serial number LRMPL1-1710018, lot number 04619100). The supernatant was transferred into a fresh tube and the bicinchoninic acid (BCA) assay (Thermo Scientific, 23223) was used to measure the protein concentration. For cytoplasmic samples depleted of nuclei and mitochondria for PISA (below), samples were homogenized in 1.1 mL of KPBS (130 mM KCl, 10 mM KH_2_PO_4_, pH 7.3). Minced tissue was homogenized twice on level 2 for 1 min each with 1 min on ice in between. Homogenate was transferred to a 1.5 ml Eppendorf tube. The glass tube was rinsed with 0.3 ml of KPBS +PI and added to a 1.5 ml Eppendorf tube. Samples were spun twice at 600 xg at 4°C for 5 min with supernatant moved to a fresh Eppendorf tube each time to remove the nuclear fraction and debris. Samples were then centrifuged at 7,000 xg at 4°C for 10 min to remove mitochondria.

For PISA analysis, cytoplasmic samples depleted of nuclei and mitochondria were diluted in half to 1 mg/ml with PISA lysis buffer (2 mM MgCl2, 1% CA630, PBS (Corning 21-031-CV)). Lysates were incubated on ice for 30 min and subdivided into 4 aliquots in PCR strip tubes. One aliquot was kept on ice to use for the total protein level quantitative proteomics. The remaining three aliquots were exposed to a single temperature (52°C, 53°C, 54°C) for 3 min followed by equilibration to 25°C for 5 min. The three temperatures (52°C, 53°C, 54°C) for each sample were combined and centrifuged at 20,817 xg for 99 min at 4°C to remove aggregated protein. Supernatant (soluble fraction) was moved to a new tube. All samples were frozen on dry ice and dried down in speedvac (**Fig. S10A**).

For tandem mass tag (TMT) proteomics, 50 μg of protein (whole brain) or 12 μg (organelle depleted cytoplasm) was dried down in a speedvac (Thermo Scientific, SPD111V), resuspended in 23 μl of 1x lysis buffer (5% SDS, 50 mM TEAB, pH8.5) and digested with Trypsin/LysC (Thermo Scientific, A41007) using S-Trap micro columns (ProtiFi, C02-micro-80) as described by the manufacturer. Eluted peptides were dried in a speedvac and resuspended in 20 μl (whole brain) or 5 μl (organelle depleted cytoplasm) of 100 mM TEAB pH 8.5. The resuspended peptides were added to a tube containing TMT tag for a final protein:TMT ratio of 1:2 (ThermoFisher, #A44521). The reaction was quenched with 0.5 μl of 5% hydroxylamine and incubated for 15 min at 25°C with shaking (300 rpm). Quenched samples were combined and dried down in aliquots with 50 μg set aside as “pre-fractionation” sample. For fractionation (whole brain only), 125 μg aliquot was resuspended in 300 μl of 0.1% TFA and sonicated for 5 min. The sample was subjected to a high pH reversed phase fractionation procedure as described by manufacturer (Pierce, #84868). Fractions 2 and 8 were combined and dried down. Fractions 3-7 were kept separate and dried down. Fractions and input samples were reconstituted with 20 μl of 5% acetonitrile, 0.1% formic acid and 3.8 μl was injected per replicate.

Liquid chromatography tandem mass spectrometry (LC–MS/MS) experiments were performed in duplicate for fractionated samples and in triplicate for unfractionated samples. Samples were run in a 145 min gradient from 10% of acetonitrile (0.1% formic acid) to 70% acetonitrile, 0.1% formic acid in data dependent mode on a nanoACQUITY UltraPerformance LC (UPLC; Waters, Milford, MA, USA) coupled to an Orbitrap Fusion Lumos Tribid mass spectrometer. MS data acquisition was collected from 400 to 1400 m/z with a resolution of 120,000. MS2 Data Dependent Acquisition was acquired with a resolution of 50000 at m/z 200 with a mass injection time of 120 ms. Other acquisition methods and column setup were identical to those described previously.(61) Raw data files were processed with MaxQuant version 2.4.11.0. with reviewed entries only of mouse uniprotKB FASTA file (downloaded 08/09/2023) to identify peptide-spectrum matches (PSM) and to consolidate replicate injections.(62) Search parameters included Trypsin/P enzyme, fixed modification of carbamidomethylation on cysteine (+57 on C) and variable modifications of oxidized methionine (+16 on M), phosphorylation of serine (S), threonine (T), and tyrosine (Y) (+80 on STY) and N-terminal acetylation (+42). The files were processed with Reporter MS2 type set to TMT 16-plex (+304 on peptide N-Terminal and Lysine). Assignments were made using a 20-ppm mass tolerance for first search and 4.5 for main search for the precursor and 0.5 Da mass tolerance for the fragments with a 1% FDR set for PSM and Protein. The proteinGroups output text file was imported into Perseus (2.0.10.0) for log_2_ transformation, imputation using values based on normal distribution, and quantile normalization.(63) Additional filters included numerical value in at least 4 of the 16 samples and 2 unique peptides for protein identification. The exported table was then processed in RStudio (R version 4.3.3) using limma (3.58.1) for differential expression (or stability) testing with a linear model that had a factor for group, whose levels were the four combinations of GT (Q331K or WT) and INJ (TBI or Sham). Empirical Bayes moderated t-statistics were computed for contrasts that included four between-group comparisons (WT.TBI vs WT.Sham, Q331K.TBI vs Q331K.Sham, WT.TBI vs Q331K.TBI, and WT.Sham vs Q331K.Sham), main effects of GT (Q331K vs WT) and INJ (TBI vs Sham), and the interaction between GT and INJ. Proteins with an FDR < 0.1 were considered significantly different for these contrasts. For the PISA data processing, one Q331K sham sample was identified as an outlier based on total intensity and excluded from further analyses. Data are available via ProteomeXchange with identifier PXD068825 and 10.6019/PXD068825.

### Western blotting

For Western blot analyses, protein samples were mixed with loading buffer (Boston BioProducts, BP-111R) and boiled at 100 °C for 5 min. Gel electrophoresis was performed with the BIO-RAD PowerPac HC followed by a transfer onto polyvinylidene fluoride (PVDF; Merck Millipore, IPFL00010) membrane. The PVDF membrane was blocked for 1h at ambient temperature using Intercept Blocking Buffer (LI-COR, 927-70003) and then primary antibody (see table below) was incubated at 4°C overnight (>12 h) in PBST and secondary antibody was incubated at ambient temperature for 1 h in blocking buffer. The PVDF membrane was imaged using the LI-COR ODYSSEY infrared imager system (Serial No. ODY-2215). The target protein intensity was quantified using Image Studio software (Lite Ver 5.2).

| Antibody | Manufacturer | Catalog # | Dilution |
| --- | --- | --- | --- |
| H3 | Proteintech | 17168-1-AP | 1:1000 |
| Calr | Enzo life sciences | ADI-SPA-602 | 1:1000 |
| Cox4 | Abcam | AB14744-1005 | 1:1500 |
| GAPDH | Sigma-Aldrich | G8795 | 1:2000 |
| Rpl4 | Proteintech | 11302-1-AP | 1:2000 |
| Rps6 | Cell Signaling | 2217 | 1:500 |
| TDP43 | Encor | MCA3H8 | 1:5000 |
| Anti-Mouse IgG 680 | LICORbio | 926-68072 | 1:10,000 |
| Anti-Rabbit IgG 800 | LICORbio | 926-32211 | 1:10,000 |

### Immunostaining

Mice were euthanized and transcardially perfused with PBS and 4% paraformaldehyde (PFA; Fisher Scientific, AAA1131336) in 1X PBS, pH 7.4. Mice were decapitated, and the scalp and skull carefully removed to harvest the intact brain. All brain samples were immersed in 4% PFA solution for 24h at 4°C for post-fixation. A brain matrix (CellPoint Scientific, Alto Acrylic 1mm Mouse Brain Coronal) was used to pre-section the brains into anterior, medial, posterior and cerebellar sections, which were embedded in paraffin by the Morphology Core (UMass Chan Medical School). Coronal sections of 7 μm thickness were prepared on a Leica RM2125 RTS and mounted onto charged microscope slides (Globe Scientific Cat#1354W). Prior to immunostaining, tissues were deparaffinized with xylenes (Fisher Scientific, X3P-1GAL) and rehydrated in 100% and 70% ethanol (Fisher Scientific, 04355223EA). Tissues were subjected to antigen retrieval using 10mM Sodium Citrate (Fisher Scientific, S25545) containing 0.05% Tween 20 (Fisher Scientific, BP337-500) at pH 6.0, heated to boiling, and then maintained at 95°C in a steamer (Black&Decker Cat# HS1050) for 30 minutes. Tissues were blocked using 1% bovine serum albumin (BSA; Fisher Scientific, BP1600-100) and 3% horse serum (Gibco, 26050-070) in PBS for 1h at ambient temperature. Tissues were probed with rabbit anti-TDP-43 (1:250 Proteintech, 10782-2-AP) and chicken anti-NeuN (Millipore Sigma, ABN91) antibodies at 4°C overnight. Tissues were incubated at ambient temperature for 1h with fluorophore-conjugated secondary antibodies (Jackson ImmunoResearch Laboratories) at 1:500 and counterstained with DAPI. Prolong Gold antifade reagent (Invitrogen, P36930) with a refractive index of 1.46 was used to mount coverslips onto the tissues.

### Confocal Microscopy

TDP-43/NeuN/DAPI immunofluorescence images were obtained using a Leica TCS SP5II (S/N: 5100001537, 11888906 BZ:01) confocal microscope. A 40x objective lens with oil was used to acquire images. Leica Application Suite Advanced Fluorescence (LAS AF) software was used for image acquisition as follows: imaging mode: XYZ; Sequential imaging: on; Format: 512x512; Pinhole: on; Speed: 400 Hz; Line average: 1; Frame average: 3, Accu for Line average: 1; Accu for Frame average: 1; Z-stack mode: Z-wide; Sequential scan mode: between frames, number of steps: 3; Z-step size: ∼0.5 μm. Using these settings, images were collected in the following order: ipsilateral cortex, ipsilateral amygdala, ipsilateral hypothalamus, contralateral hypothalamus, contralateral amygdala and contralateral cortex.

### TDP-43 localization analyses

TDP-43 nuclear-to-cytoplasmic (N/C) ratio analysis was performed with 40x confocal images described above on a per image basis with a custom macro generated with Fiji (**Additional file 2).** (64) DAPI signal was used to define the area of nuclear TDP-43 fluorescence and the cytoplasm was defined as any area outside of the DAPI-defined region. Mean TDP-43 fluorescence intensity was measured for the nuclear and cytoplasmic regions, and the N/C ratio was calculated by dividing the mean nuclear TDP-43 signal by the mean cytoplasmic TDP-43 signal.

TDP-43 N/C ratio analysis was also performed on a per-cell basis using a customized pipeline that was developed in CellProfiler(65) for calculation of the percentage of cells with increased cytoplasmic TDP-43 **(Additional file 3)**. DAPI signal was used to identify the nuclear region, which was then expanded by 8 pixels to capture a contiguous cytoplasmic region. The mean TDP-43 fluorescence intensity was measured for the defined nuclear and cytoplasmic regions and the N/C ratio was calculated as described above for the per image analysis. Cells with an N/C ratio <5.72 were classified as having cytoplasmic TDP-43. This threshold was determined using the average N/C ratio of each field of view (FOV) in the WT sham condition (representing a condition that did not exhibit robust cytoplasmic TDP-43) minus one standard deviation. Within a FOV, the number of cells with an N/C ratio <5.72 was then divided by the total number of cells, multiplied by 100 to obtain the percentage cells with cytoplasmic TDP-43 for each condition.

The CellProfiler pipeline above was also used to measure the distribution of TDP-43 signal within the nucleus and to quantify the number of cells with a “ring-like” TDP-43 staining appearance. The nucleus, as defined by DAPI, was divided into three concentric regions using the Measure Object Intensity Distribution module in CellProfiler and setting the number of bins to three. The rings are therefore proportional to the size of each nucleus and nuclear area is accounted for in the pipeline. For example, the outer ring represented the outer 1/3 of the nucleus, regardless of nuclear size. TDP-43 fluorescent signal was measured within each of the three nuclear ring-regions. Cells exhibiting ring-like TDP-43 staining were defined as those with >56% of the total nuclear TDP-43 signal in the outer region (i.e., at the boundary of the nuclear envelope). This threshold was determined by calculating the average percent of total nuclear TDP-43 signal present in the outer region of the nucleus (the outermost concentric ring) in all WT sham cells plus one standard deviation. Within a FOV, the percentage of cells with >56% nuclear TDP-43 signal in the outer nucleus out of the total cells per each condition was determined and used for analysis.

The outcomes of the TDP-43 localization analyses above were combined to generate a measure of cells with altered TDP-43 localization, which included cells that exhibited TDP-43 ring-like and/or cytoplasmic TDP-43 according to the following parameters:

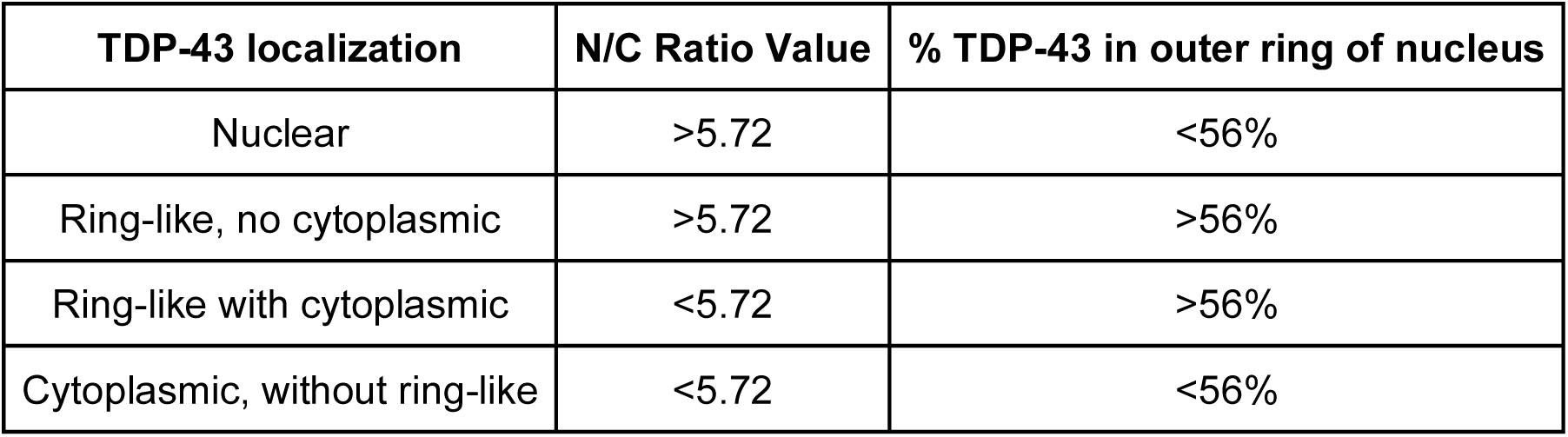

| TDP-43 localization | N/C Ratio Value | % TDP-43 in outer ring of nucleus |
| --- | --- | --- |
| Nuclear | >5.72 | <56% |
| Ring-like, no cytoplasmic | >5.72 | >56% |
| Ring-like with cytoplasmic | <5.72 | >56% |
| Cytoplasmic, without ring-like | <5.72 | <56% |

The total number of cells that demonstrated each of the four TDP-43 distributions from the above table (nuclear, ring-like +/- cytoplasmic and cytoplasmic) was summed then divided by the total cells per FOV in each condition and multiplied by 100 to calculate the percent of cells with altered TDP-43 localization per mouse. To visualize the overall effect of genotype and TBI on altered TDP-43 localization, the percent of cells with each TDP-43 distribution was averaged across all mice in each condition by FOV and plotted in a stacked bar graph in **Fig. 4**.

### Reverse transcription polymerase chain reaction (RT-PCR)

RNA was isolated as described above and cDNA was synthesized using iScript Reverse Transcription Supermix (Bio-Rad, 9170-8840) according to the manufacturer’s instructions. RT-PCR was performed on mouse tissue using 2x Taq RED Master Mix (APExBio Cat. No. K1034) with the primers listed (**Supp. Table V**) with a touchdown PCR according to the manufacturer’s instructions. Products were separated on 3.5% agarose gels using the SYBR Safe DNA gel stain (Invitrogen) and imaging was performed using the Bio-Rad Chemidoc software. Quantification of the different isoforms was performed with ImageStudio software (LICORbio), using manual settings to analyze the densitometry of each band. The intensity ratios between products with included and excluded exons were analyzed for samples from a minimum of four individual mice per group.

### Structural analysis

The mm39 reference genome in Integrative Genomics Viewer, with its default RefSeq Curated gene annotation track, was used to identify exons in *Ube3c*. The splice variant, with the corresponding exon removed from the amino acid sequence, was modeled in AlphaFold 3.(66) Structural alignments of the models were performed in PyMOL (Molecular Graphics System, Version 3.1.3.1, Schrödinger) by minimizing the root-mean-square deviation (RMSD) between superimposed structures. The resulting aligned structures were visualized in PyMOL.

### Ribosomal Profiling of cytoplasmic fractions

Ribosomal profiling was performed as previously described (67) on cytoplasmic fractions. A 20% to 50% sucrose gradient was prepared using a Gradient Master 107 (BioComp) in 14 x 89 mm polypropylene ultracentrifuge tubes (Beckman 331372). A total of 6 mL of 20% sucrose solution in 10 mM HEPES pH7.4, 150 mM NaCl, 5 mM MgCl2 with 100 μg/ml of cycloheximide, was added and overlayed with 50% sucrose solution made using the same buffer conditions. The top of the sucrose gradient (300 μl) was removed and replaced with 300 μl of sample (800 μg) containing 100 μg/ml of cycloheximide and 40 U/ml of RNaseIN. Samples were centrifuged at 197,000 x g for 2h at 4°C using a Beckman SW41 Ti rotor. Samples were loaded on a Density Gradient Fractionation system (Brandel), pushed with 60% sucrose and 0.72 ml fractions were collected with real-time UV monitoring at 254 nm using an ISCO UA-6 UV detector. Protein was precipitated from fractions using cold acetone. Protein pellets were collected by centrifugation at 13,000 x g for 10 min at 4°C, resuspend in 100 μl of SDS loading buffer, and heated for 5 min at 90°C.

## RESULTS

### Mutant TDP-43 mice exhibit heightened neurological deficits and delayed recovery following TBI

To examine the effects of TDP-43 dysfunction in the context of CNS challenge, we administered a mild TBI using a weight-drop device (**Fig. 1A**) to mice harboring a knock-in ALS-linked Q331K mutation in TDP-43.(44) TDP-43 Q331K mice (hereon referred to as Q331K mice) exhibit phenotypes consistent with frontotemporal dementia (FTD), but without motor deficits or motor neuron degeneration that is associated with ALS.(44) This mild TBI model has been described by us previously.(39, 40, 43, 51, 52, 68) Here, a free-falling 50g weight was dropped from a height of 15cm and used to deliver multiple impacts to the mouse’s exposed but intact skull within a single surgical session (**Fig. 1A,B**). Sham animals underwent the same surgery as TBI animals to retract the skin from the skull but without any impact.

TBI studies included four main groups of mice at 16-weeks of age: Q331K and WT mice, each with sham or TBI surgery. Neurological evaluations of motor function, alertness and physiological behavior pre- and post-TBI were accomplished using a set of 10 tasks.(53) The results of these tests are reported as a Neurological Severity Score (NSS), which is used to assess potential neurological deficits following TBI in rodent models, where a higher score reflects worse neurological deficits.(53). Consistent with our previous reports,(39, 43, 68) WT mice exhibited worse NSS scores after TBI compared to their sham counterparts at 2 h post-TBI. Notably, Q331K mice exhibited significantly worse NSS scores during the acute phase post-TBI (i.e., 2h, 24h, and 48h post-TBI) compared to their WT counterparts, demonstrating heightened vulnerability and delayed recovery to brain trauma **(Fig. 1C)**. All groups returned to baseline by one-week post-TBI, as expected for a mild-TBI paradigm.(43)

Mice were also evaluated for long-term deficits resulting from TBI using the cued and contextual fear conditioning (CCFC) test, an assessment of learning and memory.(69, 70) The CCFC test consisted of four days, with habituation on day 1 (**Fig. 1D**), conditioning with a paired tone and foot shock on day 2 (**Fig. 1E**), testing recall of the foot shock in response to the tone (i.e., cue) on day 3 (**Fig. 1F**), and testing recall of the CCFC chamber (i.e., context) on day 4 (**Fig. 1G**). An animal’s freezing behavior on days 3 and 4 reflects its capacity for learning and memory, where attenuated freezing is indicative of deficits in learning and memory.(69, 70) WT and Q331K mice were randomly assigned to group 1 or group 2, and both groups were subjected to the CCFC test before surgery. Group 1 and group 2 animals were then, respectively, administered sham or TBI surgery and assessed using the CCFC test again 5-months post-surgery. Linear mixed effect analyses were then used to determine potential differences between groups and over time (i.e., between pre- and post-surgery) for the CCFC test. This uncovered significant effects for the conditioning (Day 2) and cue-induced freezing response (Day 3). For conditioning, significant group (P=0.010; numerator degrees of freedom or df=3, denominator df=107.770, F=3.991) effects were detected, but not significant time (P=0.139; df=1, denominator df=107.773, F=2.220) or group x time interaction (P = 0.724; numerator df=3, denominator df=107.770, F=0.441) effects. On pair-wise testing, one of the Q331K cohorts was found to exhibit altered freezing behavior during conditioning compared to one of the WT cohorts and also to the other Q33K1 cohort (**Fig. 1E**). As mice were randomly assigned to groups 1 and 2, the reason for the latter difference in pre-surgery mice is unclear. For cue-induced freezing response, significant group (P = 0.016; numerator df=3, denominator df=107.938, F=3.595) and time (P <0.001; numerator df=1, denominator df=107.955, F=14.651) effects were detected, but not significant group x time interaction (P = 0.298; numerator df=3, denominator df=107.938, F=1.243) effects. On pair-wise testing, both Q331K cohorts exhibited attenuated cue-induced freezing on day 3 relative to WT animals, where comparisons with the WT-2 cohort reached significance (**Fig. 1F**). These outcomes are consistent with Q331K mice exhibiting cognitive deficits prior to surgery, as shown for other cognition tests described by White et al.(44)

Both WT sham and WT TBI groups showed differences in the cue-induced freezing response at 5 months post-surgery versus the pre-surgery timepoint (**Fig. 1F**). The reduced freezing behavior in WT sham animals 5 months post-surgery may stem from aging and/or adverse effects related to the sham surgery. Conversely, Q331K mice showed no difference in the cue-induced freezing response at 5 months when compared to pre-surgery, when they already exhibited cognitive deficits. Therefore, there may be a ceiling effect with respect to cognitive dysfunction in Q331K mice, given that Q331K mice exhibit similar cue-induced freezing behavior pre- and post-surgery, regardless of surgery-type or age (**Fig. 1F**). Finally, there were no significant differences in context-induced fear behavior due to genotype or surgery (**Fig. 1G**). Taken together, this mild-TBI paradigm appears to exert minimal long-term effects on learning and memory, whereas mutant TDP-43 expression may be sufficient to cause deficits in the amygdala, the brain region that influences cue-induced behavior more so than others such as the hippocampus.(69, 70)

### Proteostasis-related genes become upregulated in mutant TDP-43 mouse brain after TBI

To investigate the gene expression changes associated with enhanced neurological severity scores and delayed recovery in Q331K mice (**Fig. 1C**), bulk RNA sequencing (RNAseq) was pursued using mouse whole brain lysate near the impact site (**Fig. 1A**) at the 24h post-TBI timepoint. This analysis focused on four groups: Q331K and WT mice, each with sham or TBI surgery. Of the ∼25,000 genes that remained after filtering out genes with very low expression, our analysis identified 228 that were differentially expressed between the Q331K TBI and WT TBI mice (**Fig. 2A, Supp. Table I**).

**Figure 2.**
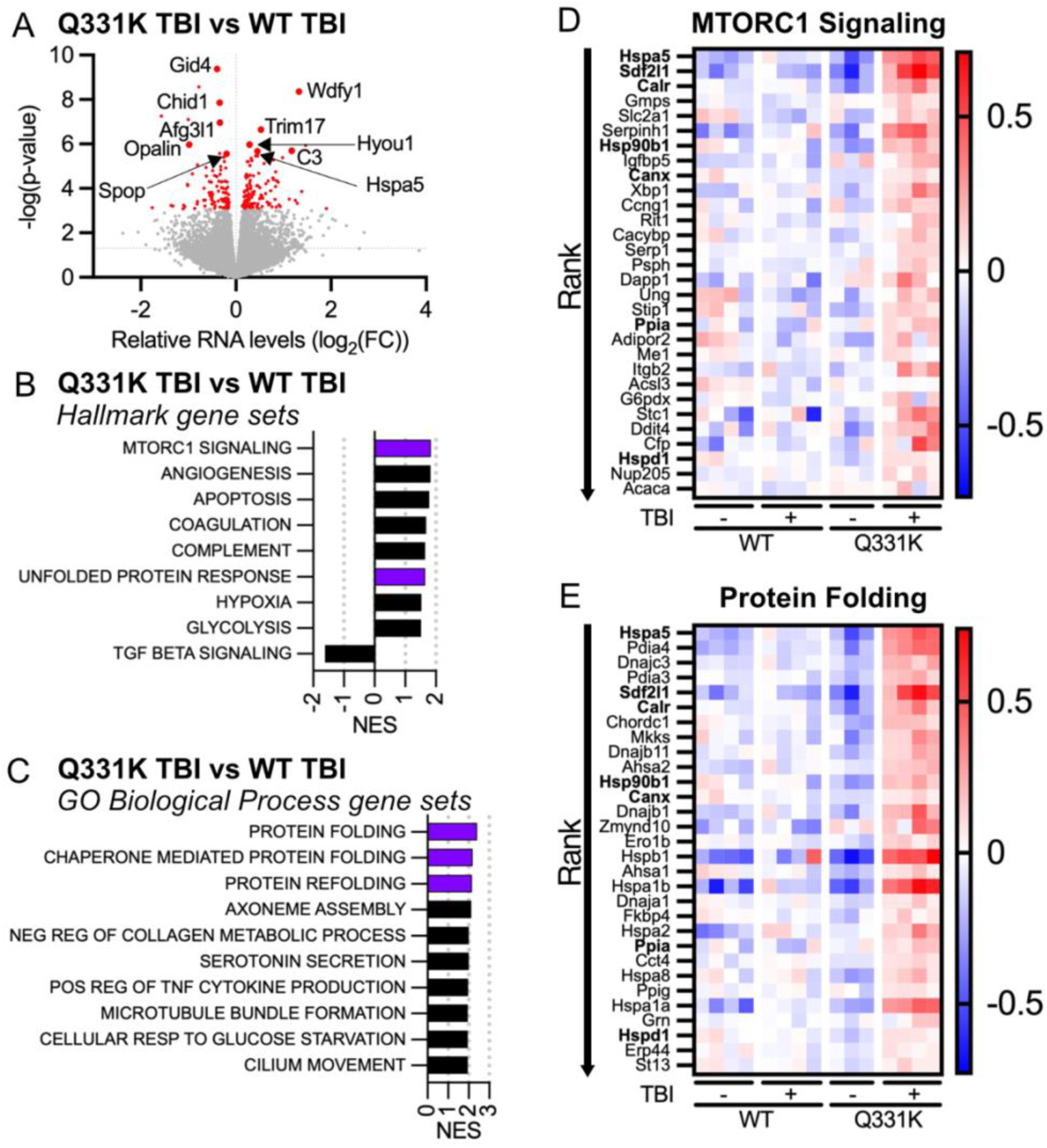
Upregulation of MTORC1 and protein folding genes in Q331K versus WT mouse brain after TBI. **(A)** Volcano plot of RNAseq data comparing Q331K and WT mice, each with either TBI or sham surgery. Differentially expressed genes (DEGs) with FDR <0.1 are shown in red. Top up- and down-regulated annotated genes are labeled. **(B,C)** Top GSEA gene set hits (both up and down regulated, as applicable) for Q331K TBI relative to WT TBI using the Molecular Signatures Database (MSigDB) hallmark **(B)** and Gene Ontology Biological Process **(C)** collections. Purple bars denote gene sets associated with proteostasis. **(D,E)** Heatmaps showing the top 30 ranked genes in the hallmark gene set “MTORC1 signaling” **(D)** and the Gene Ontology Biological Process gene set “protein folding” **(E)**. The log_2_(fold change) relative to the mean expression is shown for each sample. All pathway hits meet cutoff criteria of FDR <0.05 and |NES|>1.5. Genes in both “MTORC1 signaling” and “protein folding” gene sets are indicated in bold.

Gene Set Enrichment Analysis(59, 60) was performed using the hallmark and Gene Ontology Biological Process (GOBP) gene set collections from the Molecular Signatures Database (MSigDB).(56–58) The hallmark collection includes 50 gene sets, with clearly defined gene expression, as a high-level representation of a compilation of other gene sets in MSigDB, whereas GOBP represents >7000 gene sets capturing biological processes. A normalized enrichment score (|NES|>1.5) and an adjusted p-value cutoff (FDR<0.05) were used to determine significantly altered pathways between Q331K TBI and WT TBI mice. NES reflects the overrepresentation of a gene set at the upper (positive value) or lower (negative value) end of a ranked gene list. GSEA analysis of the hallmark collection identified “mammalian target of rapamycin complex 1 (MTORC1) signaling” as the top upregulated gene set (**Fig. 2B**; **Supp. Table II**). MTORC1 signaling is involved with multiple cellular processes, including protein synthesis and autophagy.(71) Protein folding was associated with the top 3 gene sets from GOBP identified in the GSEA analysis (**Fig. 2C**). The relative expression for the top 30 ranked genes in the “MTORC1 signaling” (**Fig. 2D**) and “protein folding” (**Fig. 2E**) pathways are shown in a heatmap representation, highlighting striking differences in gene expression between Q331K TBI mice relative to the other three cohorts. Seven genes overlap between the top ranked list for MTORC1 signaling and protein folding, with *Hspa5*, encoding the endoplasmic reticulum (ER)-associated chaperone heat shock 70 kDa protein 5, as the top ranked gene in both pathways (**Fig. 2D-E**).

To further examine differentially expressed genes between Q331K and WT mice as a function of TBI, we compared significant transcriptional changes in Q331K TBI versus Q331K sham mouse brain lysates (**Fig. S2A and Supp. Table I**) with those in WT TBI versus WT sham (**Fig. S2B and Supp. Table I)** by plotting directional p-values (**Fig. 3A**), in which -log_10_(p-value) – a measure of significance – is multiplied by the sign of the log_2_(fold-change), so that negative scores reflect a decrease in gene expression for a specific comparison and positive scores reflect an increase in expression. In this plot, genes that significantly change in expression in the Q331K TBI versus Q331K sham comparison (sections shaded in pink) can be visualized along the x-axis, whereas the y-axis highlights genes that significantly change in expression in WT TBI versus WT sham mice (sections shaded in blue). Sections shaded in grey highlight genes that are differentially expressed as a function of TBI, irrespective of genotype (**Fig. 3A**), and include 63 genes with FDR<0.1 in both comparisons (**Fig. 3B, Supp. Table I**). GSEA hallmark pathway analysis revealed “TNFa signaling via NFkB” as the top ranked, upregulated pathway in both Q331K (**Fig. 3C and Fig. S2C**) and WT (**Fig. 3D and Fig. S2D**) brain lysates post-TBI relative to their sham counterparts (**Supp Table II**). This is not surprising, given that inflammation and changes in innate immunity-related processes are well-documented following brain trauma.(72) Pathway changes that were specific to the WT TBI versus WT sham comparison indicate a downregulation of metabolic and bioenergetic processes that occur in the mitochondria, such as “oxidative phosphorylation” and “NADH dehydrogenase complex assembly” (**Fig. 3D,E and Fig. S2E,F**). TBI is known to affect mitochondrial bioenergetics, where TBI induces mitochondrial damage and results in reduced energy production.(73) Although these pathways were not significantly altered between Q331K TBI and Q331K sham brain lysates, expression of genes in these pathways were variable and potentially dysregulated in Q331K mice under both TBI and sham conditions, an observation that may reflect transient and adverse effects of mutant TDP-43 on mitochondria reported by others.(74)

**Figure 3.**
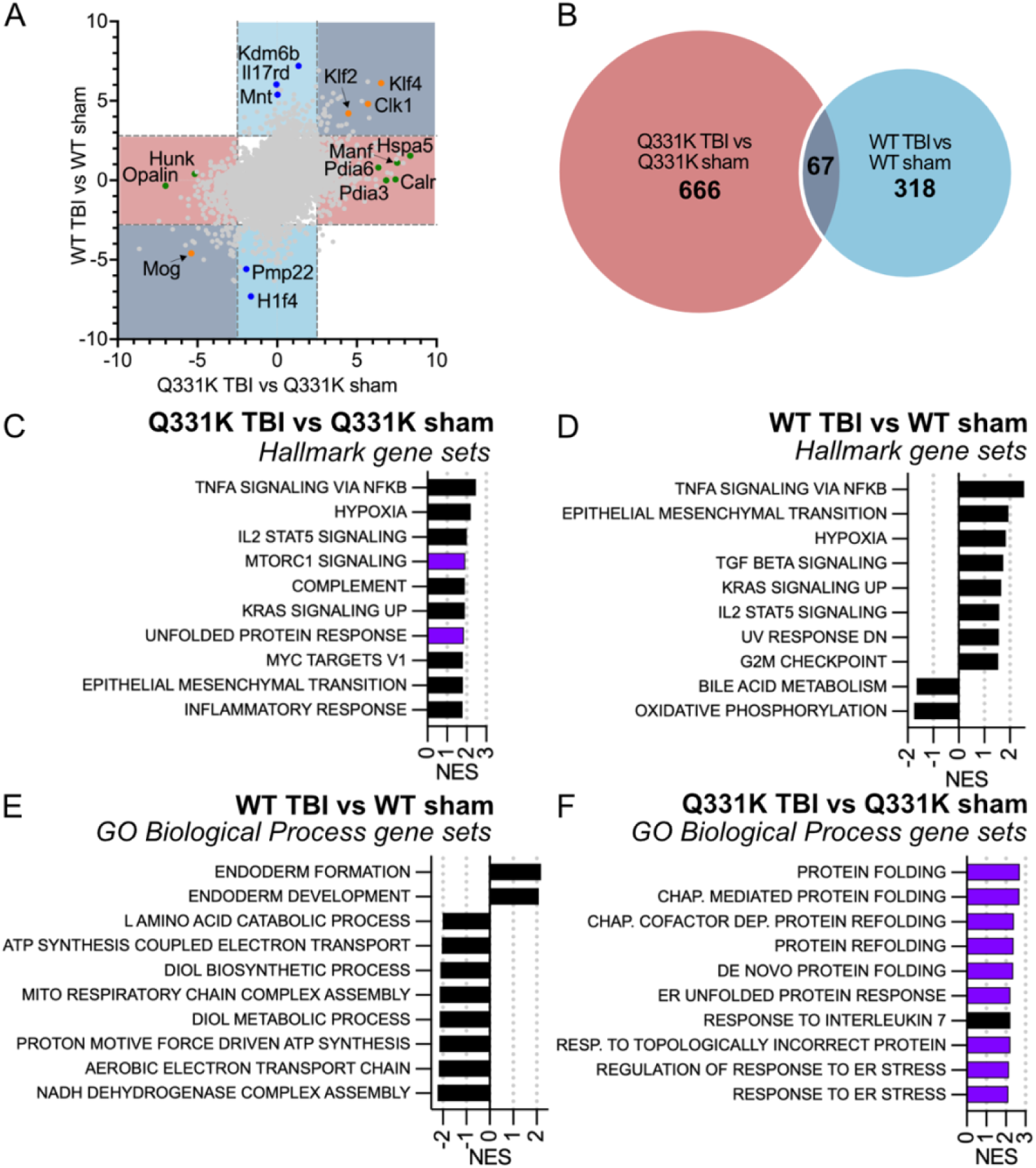
Differentially expressed genes in WT and Q331K mouse brain as function of surgery. **(A)** Plot of directional p-values (-log_10_(p-value)*sign of log_2_(fold change)) from transcriptomics datasets for the WT TBI versus WT sham comparison plotted against the Q331K TBI versus Q331K sham comparison. The sign of the score reflects the direction of the fold change. Orange dots denote genes with expression changes specifically due to TBI, irrespective of genotype. Blue and green dots denote genes with expression changes specifically in WT and Q331K mice, respectively, after TBI. Color-shaded sections indicate unique and overlapping gene changes, as illustrated in B. Dotted lines indicate FDR=0.1 **(B)** Venn diagram illustrating minimal gene overlap between Q331K and WT mice subjected to TBI using FDR <0.1. **(C-F)** Top GSEA gene set hits from the RNAseq analysis for Q331K TBI versus Q331K sham conditions using the hallmark collection **(C)**, WT TBI versus WT sham conditions using the hallmark collection **(D)**, WT TBI versus WT sham conditions using the Gene Ontology Biological Process collection **(E)** and Q331K TBI versus Q331K sham conditions using the Gene Ontology Biological Process collection **(F)**. All gene set hits meet cutoff criteria of FDR <0.05 and |NES|>1.5. Purple bars denote gene sets that are associated with proteostasis.

Strikingly, nine out of the top ten GOBP gene sets emerging from the Q331K TBI versus Q331K sham comparison were proteostasis-related, including protein folding and response to ER-stress (**Fig. 3F**), which were not enriched gene sets in the WT TBI versus WT sham comparison (**Fig. 3D, Supp Table II**). Further, proteostasis-related genes *Hspa5*, mesencephalic astrocyte-derived neurotrophic factor (*Manf*), calreticulin (*Calr*) and protein disulfide isomerase family A, members 3 (*Pdia3*) and 6 (*Pdia6*) were significantly increased in Q331K TBI versus Q331K sham mice (**Fig. 3A**; pink-shaded region). Proteostasis-related gene expression changes did not persist long-term post-TBI, as determined by a separate RNA-seq analysis on whole brain mouse lysates 7-months post-TBI or sham surgery, for both WT and Q331K mice. In fact, there were no significant gene expression changes (at cutoff FDR < 0.1) due to TBI for either genotype (i.e., Q331K TBI versus Q331K sham and WT TBI versus WT sham; **Supp Table I**).

Considering the upregulation of transcripts involved in protein folding and ER-stress in Q331K mice 24h post-TBI compared to the other three cohorts, we assessed protein level changes with quantitative proteomics analysis of mouse brain lysates labeled with tandem mass tags (TMTs) from the four cohorts. Other than TDP-43, no significantly upregulated proteins were detected (**Fig. S3A, Supp Table III**). In fact, TDP-43 upregulation was observed in Q331K versus WT comparisons irrespective of surgery type, at both mRNA and protein levels (**Fig. S3C,D**), consistent with a loss of TDP-43 autoregulation due to the Q331K mutation.(44) Despite a significant upregulation of *Hspa5* at the mRNA level in Q331K TBI mice, differential expression of Hspa5 protein was not observed across the four mouse cohorts (**Fig. S3B)**. We additionally probed for isoforms of active X-box binding protein 1 (Xbp1; **Fig. 2D**), which becomes spliced (Xbp1s) and activated under conditions of ER-stress (**Fig. 3F**),(75) by Western blot analysis. Xbp1s was not detected in samples from any mouse cohort and levels of unspliced Xbp1 (Xbp1u) were unchanged across conditions (**Fig. S3E**). The apparent lack of upregulation at the protein level may be due to deficits in protein synthesis or because 24h post-TBI is too early a timepoint for detection of newly translated protein. Nonetheless, the transcriptomics data clearly indicate a robust upregulation of proteostasis-related pathways at the mRNA level that is dependent on both TBI and expression of mutant TDP-43.

To gain insight into the relevance of our TBI model to neurodegenerative diseases such as ALS and FTLD-TDP, we compared the GSEA hallmark gene set analyses of the TBI versus sham conditions for both WT and Q331K mice with recent transcriptomic analyses of human ALS post-mortem spinal cord and FTLD-TDP brain tissues.(45, 46) Significantly upregulated gene sets in FTLD-TDP and/or ALS CNS tissues, such as “TNFa signaling via NFkB” and “hypoxia”, were also significantly upregulated in our TBI model, irrespective of genotype (**Fig. S4**). Notably, several significantly upregulated gene sets in FTLD-TDP and/or ALS CNS post-mortem tissues were only observed in Q331K mice after TBI, but not WT mice after TBI, including “MTORC1 signaling”, “unfolded protein response” and “inflammatory response” (**Fig. S4**). Similarly, GO terms related to cytoplasmic translation and innate immunity emerged from a single nucleus sequencing analysis of neurons from a separate study of human post-mortem ALS and FTLD brain tissue.(76) Therefore, there appears to be a synergistic effect of mutant TDP-43 dysfunction and acute brain trauma on biological processes in the CNS that, at least partially, recapitulates neurodegenerative disease processes on the ALS/FTLD spectrum.

### Proteostasis-related factors and pathways are downregulated in mutant TDP-43 mice at baseline

We performed similar omics analyses on cohorts of WT and Q331K mice without TBI, including RNAseq on mice without surgery (i.e., naïve mice). In our view, the sham condition represents the most relevant control for TBI, as it accounts for gene expression changes that occur from the surgery itself, thereby revealing expression changes due to the impact. Including naïve mice here serves as an additional control for identifying differentially expressed genes due to expression of mutant TDP-43. For the Q331K versus WT sham comparison, only two downregulated gene sets emerged from our GSEA analysis, “protein folding” and “aerobic electron transport chain” (**Supp. Table II**). Similarly, “MTORC1 signaling”, which includes proteostasis-related genes, such as heat shock protein family chaperones *Hspd1*, *Hspe1*, and *Hspa4*, was a downregulated pathway for the Q331K versus WT naïve comparison (**Supp. Table II**). Plotting transcriptomics data with directional p-values for the Q331K versus WT sham groups against Q331K versus WT naïve groups revealed a significant downregulation of genes involved in the ubiquitin-proteosome pathway, including glucose-induced degradation protein 4 homolog (*Gid4*) and speckle-type POZ protein (*Spop*), the latter a downregulated gene that was also observed by White et al. in this Q331K model (**Fig. S5B** and **Supp. Table IV**).(44) Of note, both *Gid4* and *Spop* remain downregulated in Q331K sham and TBI mice at 7 months post-TBI relative to WT groups (**Supp. Table I, Fig. S5C,D**). Further, our quantitative proteomics analysis of Q331K versus WT sham groups uncovered a modest yet significant decrease in the large ribosomal subunit protein P1 (Rplp1), which plays a critical role in embryonic and brain development (**Fig. S3F**, **Supp. Table III**).(77) Collectively, these omics data point toward a deficit in proteostasis-related pathways in Q331K mice at baseline.

### TDP-43 exhibits altered nuclear and cytoplasmic localization following TBI

Different TBI models reportedly induce subcellular redistribution of TDP-43 from the nucleus, where the protein is predominately expressed under homeostatic conditions, to the cytoplasm.(13, 37–41) Enhanced cytoplasmic localization of TDP-43 is also well-documented in other stress paradigms in vivo and in vitro, indicating that subcellular redistribution of TDP-43 represents a physiological response to stress.(2–6, 78) Given the disparate responses between Q331K and WT mice to TBI (**Figs. 1-3**), we examined the cellular localization of TDP-43 in situ as a function of surgery and TDP-43 genotype. Mouse brain tissue sections from TBI and sham cohorts were stained with anti-TDP-43 antibody and counterstained with DAPI. Obvious changes in subcellular TDP-43 distribution were observed in the cortex, amygdala, and hypothalamus (regions defined in **Fig. 1B**) of both Q331K and WT mice (**Fig. 4**), whereas changes in TDP-43 subcellular distribution were minimal or not detected in other regions of the tissue section (not shown). Sham animals also exhibited some degree of cytoplasmic TDP-43 redistribution in these brain regions, possibly indicative of cellular “stress” caused by the sham surgery, and therefore the naïve condition was included here for further analysis.

**Figure 4.**
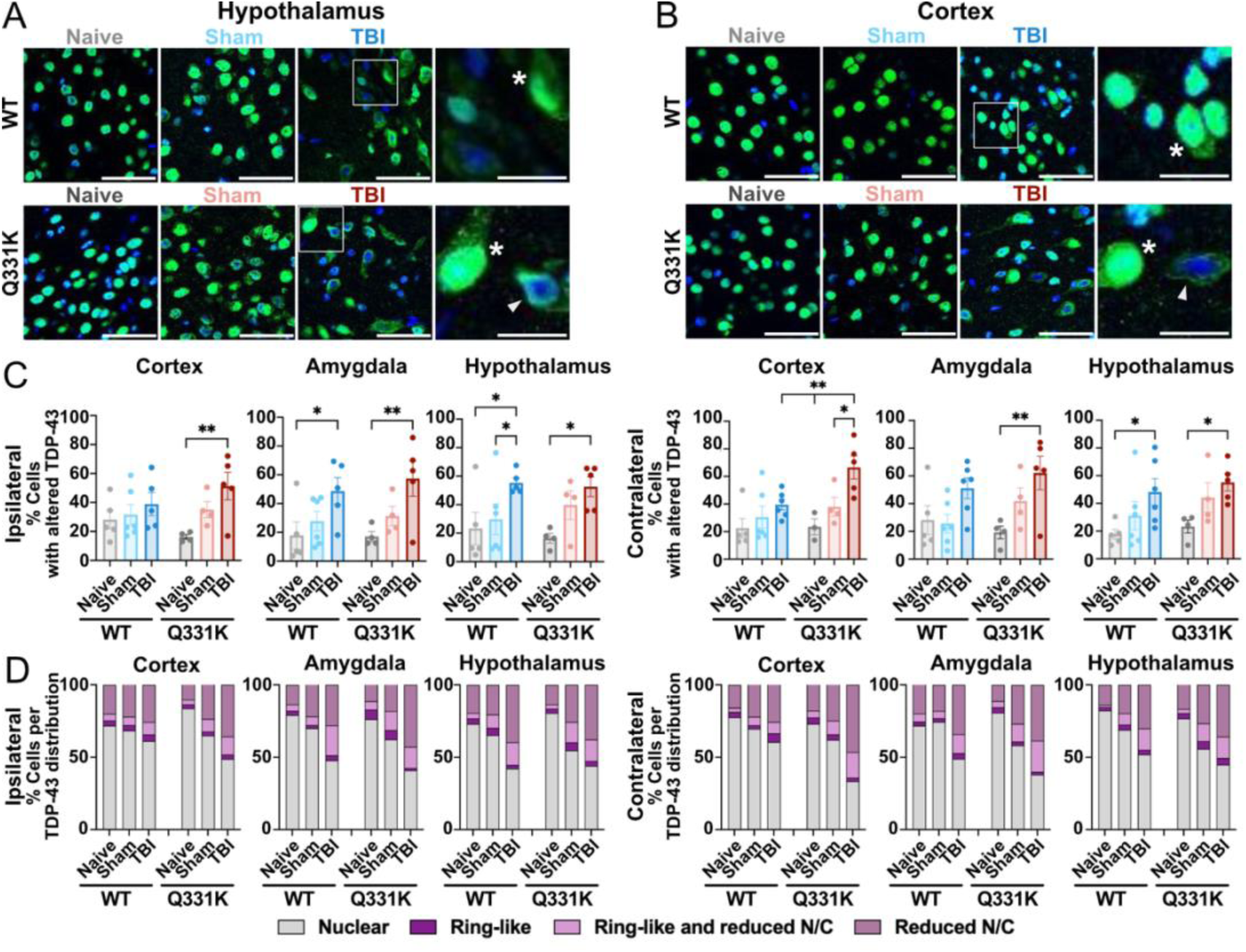
Alterations in TDP-43 localization in WT and Q331K mice as a function of surgery. **(A-B)** Confocal immunofluorescence images from WT and Q331K mice under native (i.e., no surgery) conditions or 24h post-surgery (sham and TBI). Tissues were stained for TDP-43 (green) and counterstained with DAPI (blue). Images from the hypothalamus (A) and cortex (B) are shown. Cropped insets highlight cells with cytoplasmic (asterisks) and ring-like distribution (triangles) of TDP-43 under conditions of TBI. Scale bar=50 um in the low magnification image and =20 um in the insets. **(C)** Percent of cells that exhibit altered TDP-43 localization (ring-like, reduced nuclear-to-cytoplasmic (N/C) ratio, or both) in the cortex, amygdala, or hypothalamus, either ipsilateral (left) or contralateral (right) to the impact site 24h post-surgery. (**D)** Stacked bar graphs representing the average percent of cells that exhibit nuclear TDP-43 localization (grey) or a type of altered TDP-43 localization (defined in C and color-coded according to the key) within the indicated brain region. **(C-D)** Quantification of TDP-43 localization was performed using tissues from n=4-6 mice, where each dot represents a different mouse. Statistical analysis was performed using two-way ANOVA followed by Tukey’s multiple comparisons test: *p<0.05, **p<0.01, ***p<0.001, ****p<0.0001.

To quantify TDP-43 cellular localization, confocal microscopy images were acquired within the cortex, amygdala, and hypothalamus of Q331K and WT mice in the absence of surgery (i.e., naïve) or 24h after receiving a sham or TBI surgery, for a total of six groups (**Fig. 4A,B**). First, TDP-43 nuclear-to-cytoplasmic (N/C) ratios were determined for all cells across a field of view (FOV) on a per image basis. Reductions in TDP-43 N/C in TBI versus naïve conditions were generally observed in all three brain regions for both WT and Q331K mice, although some comparisons did not reach statistical significance (**Fig. S6A**). Notably, TBI-induced reductions in TDP-43 N/C ratios occurred in both ipsilateral (i.e., TBI side) and contralateral hemispheres of the brain (**Fig. S6A**) and were prominent in the amygdala and hypothalamus. Therefore, the effects of TBI on subcellular TDP-43 localization extend beyond the site of injury in the cortex, likely because the weight-drop induces contact between the skull and brain tissue at the base of the skull (**Fig. 1A,B**). TBI-induced changes in TDP-43 subcellular localization represented an acute response to TBI, as differences in TDP-43 N/C ratios were no longer observed at 5-months post-surgery (**Fig. S7**).

As changes in TDP-43 subcellular localization were not observed in every cell, the percentage of cells exhibiting reduced TDP-43 N/C ratio relative to the naïve condition was determined from the FOVs above. Q331K TBI mice exhibited a significantly higher percentage of cells with reduced TDP-43 N/C ratio in all three brain regions relative to Q331K naïve mice for both ipsilateral and contralateral hemispheres, which was collectively a more robust change than for their WT counterparts (**Fig. S6B**). During this analysis, we also noticed a “ring-like” TDP-43 staining pattern, which appeared as dense TDP-43 signal near the outer edge of (but still within) the nucleus (**Fig. 4A**, Q331K inset) and that also occurred in cells with cytoplasmic TDP-43 staining. We developed a CellProfiler pipeline (see Methods) to quantify the percentage of cells with this phenotype and found significant ring-like TDP-43 staining in TBI versus control comparisons in all brain regions for at least one genotype (**Fig. S6C**). Ring-like TDP-43 staining was most prominent in the hypothalamus and amygdala for both Q331K TBI and WT TBI mice versus their respective controls, on both sides of the brain (**Fig. S6C**).

To assess the full extent of altered TDP-43 subcellular localization after TBI, we compiled the data above (**Fig. S6**) and determined the percentage of cells exhibiting reduced TDP-43 N/C ratio and/or ring-like TDP-43 (**Fig. 4C,D**). TBI induced more pronounced changes in TDP-43 localization in the cortical region below the impact site in Q331K versus WT mouse brain, particularly on the contralateral side (**Fig. 4C,D**). These analyses also highlighted that Q331K sham animals exhibited significant TDP-43 subcellular localization changes that were more similar to the Q331K TBI condition and were significantly different than their Q331K naïve counterparts, especially in the hypothalamus (**Fig. S6B,C**). Taken together, these data demonstrate an acute response involving the TDP-43 protein to brain injury that occurred irrespective of genotype. That Q331K mice appeared to be more sensitive to both sham and TBI surgery with respect to changes in subcellular TDP-43 localization further supports the notion that TDP-43 dysfunction leads to altered stress responses in vivo.

### Effects of TDP-43 mutation and TBI on select TDP-43 splice targets

An important nuclear function of TDP-43 involves the alternative splicing of conserved exons for numerous RNA targets,(1) which can become mis-spliced as a result of TDP-43 knock-down, overexpression and/or mutation.(79, 80) We therefore considered whether TBI-induced subcellular redistribution of TDP-43 (**Fig. 4** and **Fig. S6**) would manifest in altered splicing of TDP-43 RNA targets. To address this possibility, we probed for potential changes in splicing patterns for eight known RNA targets of TDP-43 within mouse brain lysates derived from near the TBI site (**Fig. 1A**) using RT-PCR and primer sequences that cover the exon junctions of interest (**Supp. Table V**). TDP-43 RNA targets included several that become mis-spliced due to TDP-43 misexpression, including DnaJ heat shock protein family (Hsp40) member 5 (*Dnajc5*), potassium voltage-gated channel interacting protein 2 (*Kcnip2*), protein phosphatase 3 catalytic subunit alpha (*Ppp3ca*), and translin (*Tsn*).(81–84) Splicing patterns were assessed by gel electrophoresis (**Fig. 5A**) and reported as the fraction of the mis-spliced variant versus the common or more abundant variant expressed under normal conditions (**Fig. 5B**). For *Dnajc5*, no differences in splicing of exon 5 were observed between the mouse groups studied herein. In contrast, enhanced exon exclusion most consistent with TDP-43 loss-of-function was detected for Q331K mice, irrespective of surgery, for *Kcnip2* (exon 3 only, indicated by an increase in S1), *Ppp3ca* and *Tsn*.(84) Consistent with TDP-43 gain-of-function reported in this and other Q331K mouse models,(44, 82) enhanced exclusion of exon 17b in Sortilin-1 (*Sort1*) was observed in both Q331K TBI and sham mice; in contrast, inclusion of exon 17b correlates with TDP-43 loss-of-function.(29, 81–83)

**Figure 5.**
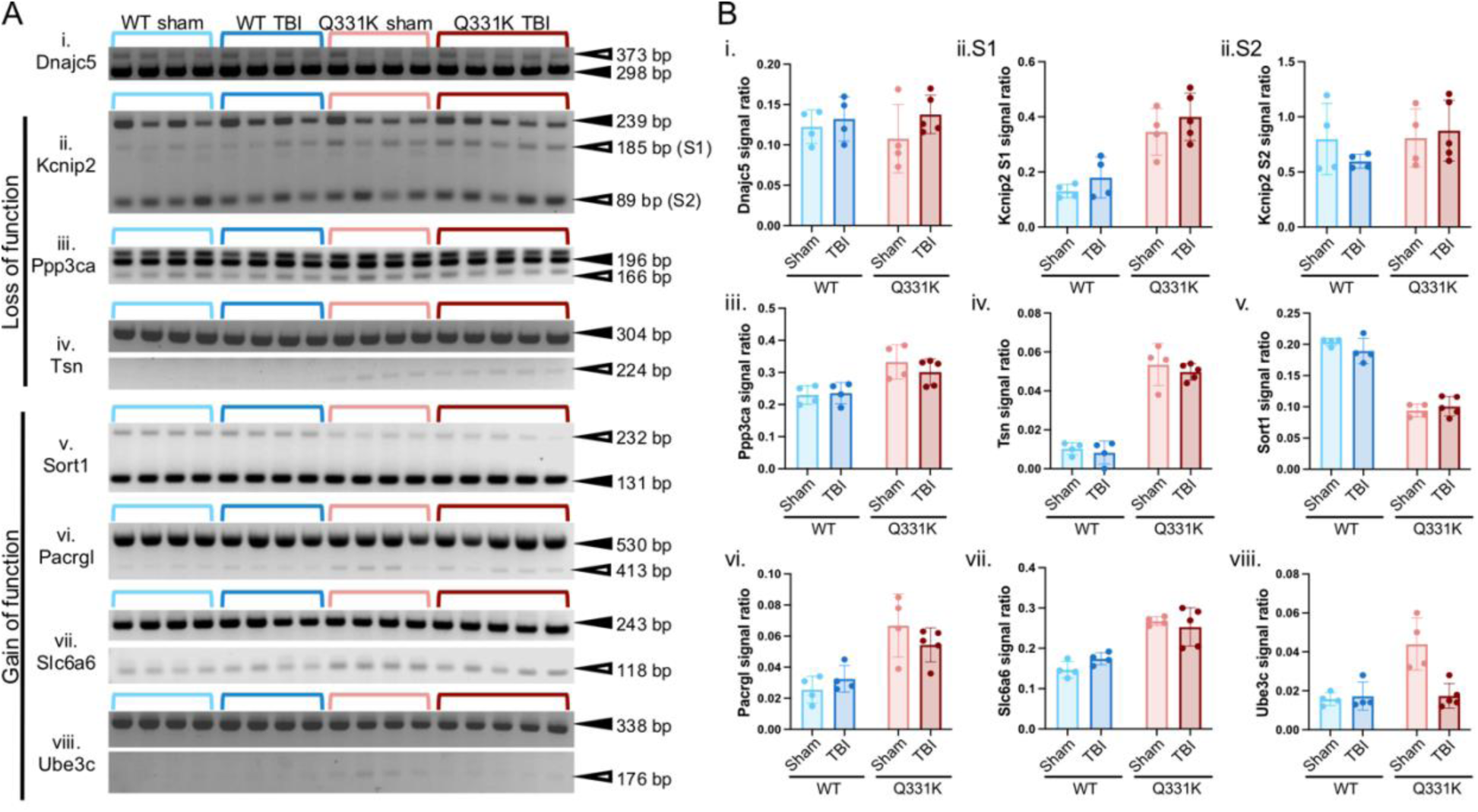
Alternative splicing patterns for TDP-43 targets in WT and Q331K mouse brain after TBI. **(A)** Splice variants of known TDP-43 RNA targets (i-vi) detected by RT-PCR analysis in WT and Q331K mice 24h after either TBI or sham surgery. The PCR primers, exons of interest and references for the primers used in this study are shown in **Supp Table VI**. Splicing patterns that are consistent with TDP-43 loss of function or gain of function are indicated. Common and mis-spliced variants are denoted by, respectively, closed and open arrowheads with the corresponding size (bp=base pairs) of the PCR product. Gels are representative of at least n=3 separate RT-PCR experiments. **(B)** Densitometry analysis of the gels in (A) was used to determine the ratio of the mis-spliced/common PCR product for each gene of interest. Each dot in the graph represents a separate mouse.

We examined additional genes with known skiptic exons caused by mutant TDP-43 gain-of-function in vivo.(80, 82) These included parkin coregulated like (*Pacrgl*), solute carrier family 6 member 6 (*Slc6a6*) and ubiquitin protein ligase E3C (*Ube3c*).(82) Skiptic exons were detected in Q331K samples for all three genes. Intriguingly, the skiptic exon in *Ube3c* was attenuated in Q331K mice after TBI, thus representing a gene-splicing event that was modulated by TBI, specifically in Q331K mice (**Fig. 5A,B**). *Ube3c* encodes an E3 ubiquitin ligase that promotes proteasomal degradation of misfolded proteins.(85) TDP-43-mediated skipping of exon 6 in *Ube3c* causes the removal of 54 amino acid residues between the ubiquitin-binding domain and the catalytic Homologous to the E6-AP Carboxyl Terminus (HECT) domain (**Fig. S8A**). While the impact of exon 6 removal on Ube3c function is unknown,(82) modeling the 3-dimensional protein structure of Ube3c in AlphaFold(66) suggests that omission of exon 6 could induce conformational perturbations in the protein N-terminal to the HECT domain (**Fig. S8B**). The effects of TBI on *Ube3c* exon 6 skipping in Q331K mice were acute, as exon skipping was no longer observed at 7-months post-TBI. Additionally, the splicing patterns for the other seven candidate TDP-43 mRNA targets were unchanged at 7-months post-TBI compared to 24h post-TBI (**Fig. S9**). Taken together, these RT-PCR results indicate that TBI-induced changes in subcellular TDP-43 localization do not strongly correlate with a loss of TDP-43 splicing function, as most of the observed splicing differences were due to TDP-43 mutation and not TBI. An exception is TDP-43 Q331K-induced skipping of exon 6 in *Ube3c*, which is attenuated during the acute phase post-TBI.

### Ribosomal proteins exhibit thermal instability in Q331K mice

A cytoplasmic shift in TDP-43 localization appears to represent a physiological response to brain trauma in mice, irrespective of TDP-43 genotype (**Fig. 4**). We hypothesized that mutations in TDP-43 could alter the nature of this response by perturbing cytoplasmic cellular processes involved in stress recovery. For example, disease-linked mutations in TDP-43 have been shown to affect the interaction of TDP-43 with cytoplasmic RNA granules as well as the biophysical properties of these granules.(11, 12, 14–16) Here, we examined protein content and protein thermostability of mouse brain lysates in both WT and Q331K mice after sham or TBI surgery using the Proteome Integral Solubility Alteration (PISA) assay (**Fig. S10A**) coupled with TMT quantitative proteomics using mass spectrometry.(86, 87) PISA capitalizes on the principle that proteins become insoluble when denatured. By extension, how a protein partitions between the soluble and insoluble fraction as a function of temperature can be used to detect changes in protein thermostability under various conditions. PISA and similar assays were historically used to elucidate protein targets of small molecules, which generally induce an increase in thermostability of their protein targets upon binding.(86, 88) We reasoned that PISA could be repurposed here for identifying thermostability shifts of cytosolic proteins and protein complexes in brain lysates as a function of TBI and TDP-43 mutation. To simplify sample processing and analysis, lysates were first depleted of large organelles. A Western blot analysis with histone H3 and cytochrome c oxidase subunit IV (CoxIV) antibodies confirmed that this protocol depleted nuclei and mitochondria, respectively, whereas anti-calreticulin confirmed that ER was retained (**Fig. S10B**).

For the PISA assay, lysates were subjected to a narrow temperature gradient (52-54°C) followed by centrifugation to remove the insoluble protein (**Fig. S10A**). The resulting soluble fraction was subjected to quantitative proteomics (**Fig. S10C**). In parallel, quantitative proteomics was performed on cytosolic lysates without heat exposure to determine total protein levels (**Fig. S10D**). Only modest differences in total protein levels between Q331K TBI and WT TBI cytosolic lysates were detected by the proteomic analysis (**Fig. S10D and Supp. Table VI)**, for which no gene sets were significantly enriched by GSEA (**Supp. Table VII**). In striking contrast, GSEA analysis of the PISA proteomics for the Q331K TBI versus WT TBI comparison revealed multiple downregulated processes, with seven of the top ten GOBP hits including terms for translation and ribosome (**Fig. 6A**). As shown in the heatmap for “cytoplasmic translation” (**Fig. 6B**), ribosomal proteins account for most of the top 30 proteins in this pathway. Proteins in the “cytoplasmic translation” pathway become elevated within the soluble fraction from WT animals after TBI compared to their sham counterparts, as demonstrated in the heatmap (**Fig. 6B**) and a box plot for the top five ranked proteins in the heatmap, all of which are ribosomal proteins (**Fig. 6C**). Conversely, these ribosomal proteins are diminished within the soluble fraction from Q331K sham mice relative to WT mice, and while they do become enhanced after TBI, their levels remain significantly below that of WT TBI lysates (**Fig. 6B,C**).

**Figure 6.**
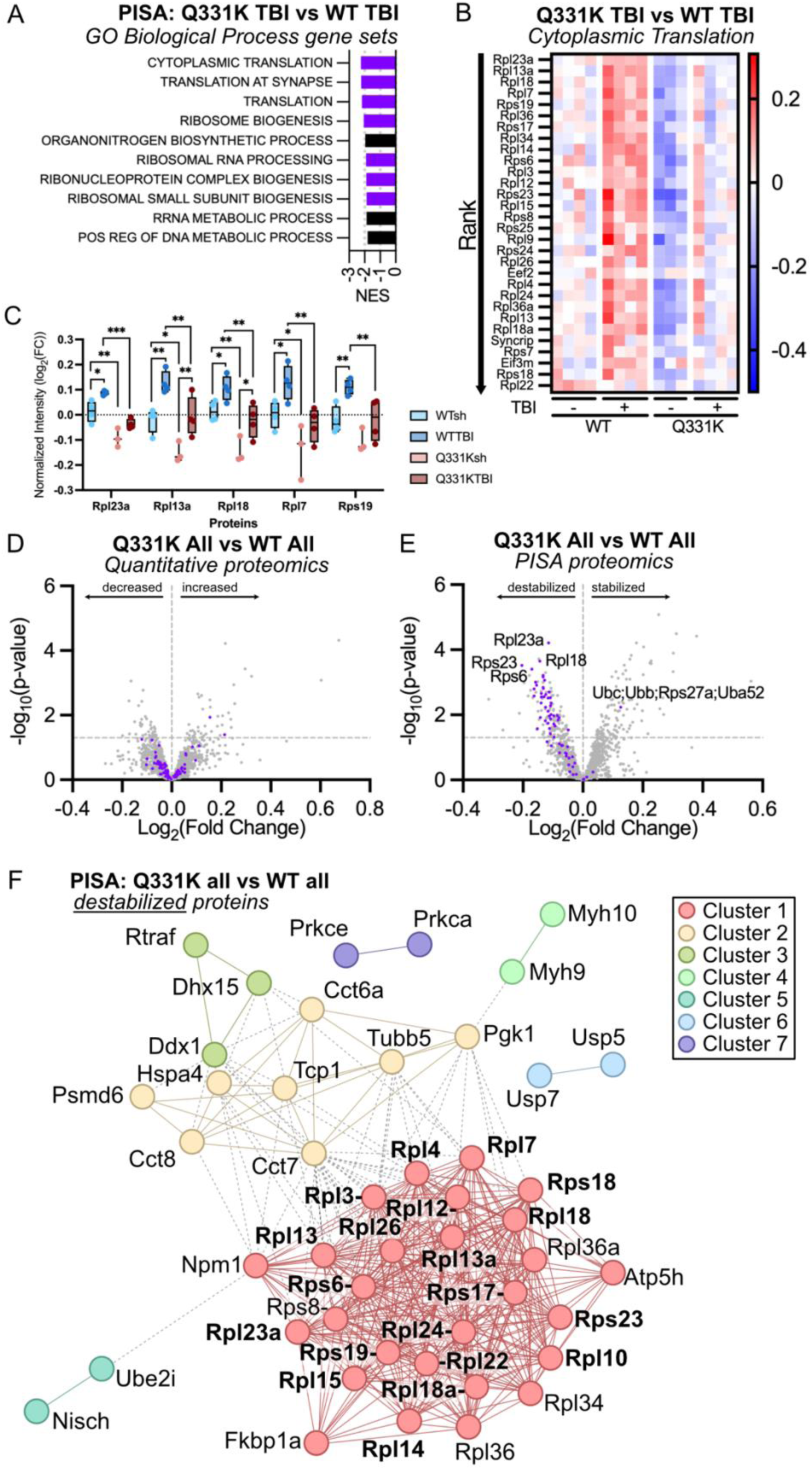
PISA analysis identifies melting temperature shifts in ribosomal and chaperonin proteins in Q331K mouse brain lysates. **(A)** Top 10 gene set hits with GSEA enrichment of thermostability changes in Q331K TBI versus WT TBI brain lysates using GOBP. Purple bars denote gene sets involved in ribosome biology. **(B)** Heatmap showing levels of soluble protein detected by PISA for the top proteins in the pathway “Cytoplasmic Translation”. The log_2_(fold change) relative to the mean expression is shown for each sample. **(C)** Box plot of top-ranking proteins in the “Cytoplasmic Translation” gene set for the indicated condition. **(D,E)** Volcano plot for the quantitative proteomics (**D**) and PISA (**E**) results for Q331K (“Q331K All”) versus WT (“WT All”), where TBI and sham conditions are combined for each genotype to assess the main effects of genotype while controlling for baseline differences related to injury. Ribosomal proteins are highlighted as purple dots. (**F**) Results of a STRING analysis using the Markov Cluster Algorithm showing significantly destabilized proteins (FDR < 0.1) in the “Q331K All” versus “WT All” comparison. Key terms included for each cluster are as follows. Cluster 1: cytoplasmic translation, Signal Recognition Particle (SRP)-dependent co-translational protein targeting to membrane, ribosome; Cluster 2: positive regulation of establishment of protein localization to telomere, association of TriC/CCT with target proteins during biosynthesis, ATP-dependent protein folding chaperone; Cluster 3, tRNA-splicing ligase complex; Cluster 4: Myosin II filament, cell shape; Cluster 5: Nisch, Ube2i; Cluster 6: synthesis of active ubiquitin: roles of E1 and E2 enzymes; Cluster 7: SHC-transforming protein 1 (SHC1) events in Erythroblastic Blast 2 (ERBB2) signaling, calcium-dependent protein kinase C activity, presynaptic cystosol. String protein-protein interaction enrichment p-value < 1.0e^-16^. Ribosomal proteins found to interact with TDP-43 in other studies are in bold within Cluster 1.(18, 91, 92)

Considering that decreased protein solubility in the PISA assay is indicative of reduced protein thermostability, our interpretation of the PISA outcomes is that ribosomal proteins and potentially ribosome complexes themselves exhibit reduced thermostability in Q331K mice relative to WT mice. Polysome profiling of cytosolic lysates confirmed that intact 80S ribosomes were indeed present for both WT and Q331K conditions (**Fig. S11A-C**). RNA fragment analysis of isolated RNA confirmed the presence of both 28S and 18S ribosomal RNA (**Fig. S11D**). Given that the effect of TDP-43 Q331K expression on ribosome stability occurred irrespective of surgery, and that moreover there were no proteins for which the genotype x injury interaction contrast was significant (FDR < 0.1) in the limma linear model, we focused on the main effect of genotype for additional proteomic analyses. This approach further verified that total levels of ribosomes in the cytosolic lysates were unchanged between TDP-43 genotypes (**Fig. 6D**), but that their thermostability was reduced in Q331K mice relative to WT mice (**Fig. 6E**). We note that Rplp1, which was reduced in the proteomics analysis of total brain lysates (**Fig. S3F**), was not quantifiable in the cytosolic proteomics because only one unique peptide was identified whereas ≥2 were required in our pipeline for protein level quantification (**Supp. Table VI**). A subsequent STRING analysis using Markov Clustering (MCL) for the 50 destabilized proteins (FDR < 0.1) revealed a major cluster of 27 proteins (Cluster 1) associated with “cytoplasmic translation” and “ribosome” (**Fig. 6F**).(89, 90) Importantly, most of the ribosomal proteins in this cluster were shown to immunoprecipitate with TDP-43 from cultured cell and/or mouse brain lysates, raising the possibility that association of mutant TDP-43 with ribosomes and/or ribosomal proteins could impact ribosome stability.(18, 91, 92) For the polysome profiling study, we noticed that the absorbance at 254nm was consistently lower for Q331K cytosolic lysates relative to that of WT within the peak where medium polysomes elute from the gradient (**Fig. S11B**). Additionally, we observed a subpopulation of TDP-43 associating with the 80S ribosome (**Fig. S11C**). These observations suggest that translation efficiency may be reduced in Q331K mouse brain relative to WT mice. “Cluster 2”, the second largest cluster revealed by STRING, includes multiple proteins involved in protein folding such as heat shock 70 kDa protein 4 (Hspa4) and four of the eight subunits that comprise the eukaryotic cytosolic T-complex protein 1 (TCP-1) ring complex or chaperonin containing TCP-1 (TRiC/CCT), including Tcp1/Cct1, Cct6a, Cct7 and Cct8. Interestingly, Cheng et al. identified these as putative TDP-43 interactors.(91) TRiC/CCT plays important roles in maintaining proteostasis and ensuring the proper folding of critical cellular proteins, lending further support to the notion that proteostasis-related processes are vulnerable and dysregulated in mutant TDP-43 mice.(93)

There were 40 proteins that exhibited enhanced stabilization in Q331K versus WT cytosolic lysates, although no cluster larger than 5 proteins was observed with STRING analyses and MCL clustering. Cluster 1 and 2 each contain 5 proteins associated with smooth muscle contraction and endocytosis, respectively (**Fig. S10G**). Interestingly, PISA detected four peptides derived from ubiquitin (**Fig. 6E**) with sequence homology shared amongst ubiquitin C, ubiquitin B, ubiquitin A-52 (Uba52) and Rps27a (**Supp. Table VI**). Uba52 and Rps27a are both ubiquitin-ribosomal fusion proteins.(94) Therefore, despite downregulation of proteostasis-related processes in Q331K mice at baseline, some compensatory mechanisms may be at play to overcome these adverse effects.

## DISCUSSION

Herein, we show that expression of dysfunctional TDP-43 leads to deficits in multiple branches of the proteostasis network (**Fig. 7**). Proteostasis deficits occur independently of overt TDP-43 overexpression, aggregation or mislocalization, which were not observed here or in prior reports describing this TDP-43 Q331K knock-in mouse model.(6, 44, 95) Analysis of our RNAseq datasets comparing Q331K and WT mice without brain trauma (i.e., naïve or sham conditions) revealed downregulation of genes involved in protein folding, including multiple heat shock proteins and chaperones, as well as genes involved in the ubiquitin proteosome pathway, such as *Gid4* and *Spop*. Exon skipping of *Ube3c*, an RNA target of TDP-43 that encodes an E3 ubiquitin ligase, was also detected in brain lysates from Q331K mice. This gain of TDP-43 mis-splicing function was initially reported in TDP-43 mouse strains with endogenous mutations in the low complexity domain, including a Q331K variant.(82) Although changes in *Ube3c* expression levels were not detected by RNAseq, AlphaFold and PyMol analyses predict that mis-splicing of *Ube3c* could alter the structure of Ube3c near the ubiquitin-binding domain. Further, Fratta et al. reported that about 15% of the exon skipping events identified in mutant TDP-43 mice occurred in genes within the ubiquitin proteosome pathway.(82) A decrease in proteosome-encoding transcripts early in life is a predictor of shortened lifespan in killifish, a vertebrate model used for aging studies.(96) Together, these observations support the notion that dysfunctional TDP-43 adversely affects the quality control arm of the proteostasis network, which in turn correlates with a decline in health and life-expectancy.

**Figure 7.**
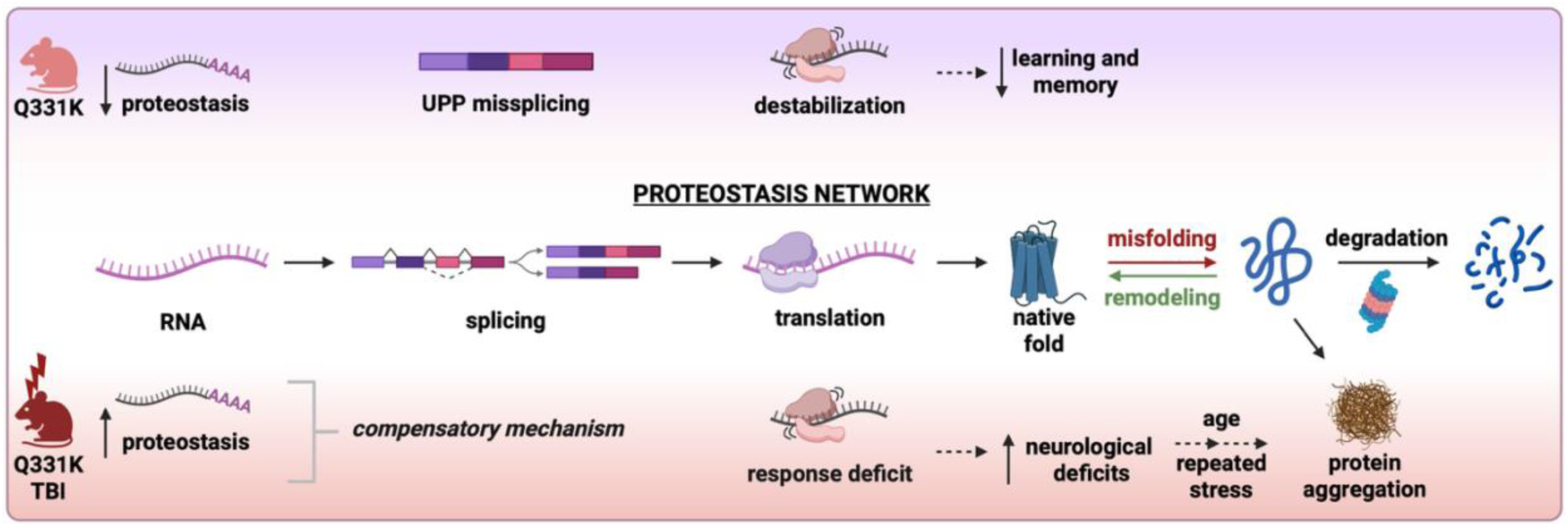
Proposed model for how the proteostasis network is affected by TDP-43 dysfunction. The proteostasis network is illustrated in the center of the diagram. Under baseline conditions (top), genes involved in proteostasis and the ubiquitin-proteasome pathway (UPP) are downregulated and mis-spliced in Q331K mouse brain relative to WT mice. In Q331K mouse brain lysates, ribosomal proteins are destabilized, which may compromise protein synthesis and thus contribute to the cognitive deficits observed in Q331K mice. In response to CNS challenge in form of a TBI (bottom), compensatory mechanisms appear to be activated in Q331K mice, including an upregulation of proteostasis-related genes at the RNA level. However, Q331K mice fail to mount a physiological response to TBI, as demonstrated by worse neurological deficits and delayed recovery after brain injury compared to their WT counterparts. We posit that destabilization of ribosomal proteins and chaperone complexes involved in proteostasis underlies the enhanced neurological deficits exhibited by Q331K mice after TBI. Although Q331K mice ultimately recover from TBI, we predict that repeated CNS challenge as a function of aging will exacerbate the phenotypes uncovered herein, and could lead to neurodegeneration and pathological TDP-43 aggregation. Model figure generated with BioRender.

PISA, a novel approach for identifying alterations in biological complexes through thermostability measurements,(86, 87) further highlighted protein folding as a vulnerable component of the proteostasis network in Q331K mice, where multiple proteins within the essential molecular chaperone TRiC/CCT were found to be destabilized. Further, 20 out of 79 core ribosomal proteins were found to exhibit reduced thermal stability through our PISA analysis.(97) These data suggest that complexes formed by ribosomal proteins, namely ribosomes, could be affected by expression of dysfunctional TDP-43 in a manner that compromises this macromolecular complex (**Fig. 7**). TDP-43 has been shown to associate with ribosomal proteins through co-immunoprecipitation and proximity labeling experiments,(18, 48, 91, 92) and with actively translating ribosomes in cultured cells.(47, 48) Ribosomal proteins have also been detected within pathological TDP-43 aggregates(48, 98) and in amyloid-positive aggregates within the aging brain.(96) In various contexts, TDP-43 expression can either repress or activate protein synthesis.(49, 50) Interestingly, the association of TDP-43 with translation-related factors can be modulated by the phase-separation behavior of TDP-43.(18) Proper phase separation of TDP-43 in vitro and in vivo requires residues 321-340, referred to as the conserved region (CR).(17–19) Deletion of the CR resulted in enhanced interactions between TDP-43 and translation factors together with heightened rates of global protein synthesis in vivo.(18) While the Q331K variant studied herein also reportedly disrupts TDP-43 phase separation,(17, 19) our polysome profiling data suggest that translation efficiency is reduced in Q331K mouse brain relative to WT brain. Therefore, there may be differential effects of the CR on translational regulation depending on how the TDP-43 sequence is modified (i.e., deletion versus single point mutation). We note that PISA was performed on organelle-depleted brain lysates in this study, such that changes to nuclear and mitochondrial proteins would not have been captured in this analysis. Indeed, it will be interesting to extend PISA for analysis of other cellular compartments and organelles in future studies.

It remains to be determined precisely how expression of mutant Q331K in mice leads to reduced thermostability of ribosomal proteins. Disease-linked mutations may perturb interactions between TDP-43 and ribosomal proteins,(91) which could induce conformational changes within the ribosome (**Fig. 7**), other complexes that include ribosomal proteins and/or free ribosomal proteins.(99) Ribosomal protein stability could also be indirectly affected by mutant TDP-43 expression. For example, altered proteostasis, activation of stress-response pathways and other stimuli can affect the post-translational modification status of ribosomal proteins, which may impact ribosome conformation and activity.(100) Intriguingly, ribosomal proteins underwent a shift toward enhanced stability in WT animals after TBI, possibly representing a mechanism that protects these proteins during cellular stress.(101) Although we do not observe changes in overall levels of ribosomal proteins in WT or Q331K mice, expression of mutant TDP-43 could influence the stoichiometry of ribosomal proteins, if not at the age of mice studied here (i.e., 16 weeks), potentially in older mice. The stoichiometry of ribosomes is reportedly compromised during aging, possibly a side-effect of impaired proteosome activity, which may lead to an accumulation of “orphan” ribosomal proteins that are prone to aggregate.(96, 101) TDP-43 dysfunction has also been linked to deficits in transporting mRNAs that encode ribosomal proteins(102) and in the stability of ribosome-encoding transcripts,(103) both of which could also impact ribosome integrity and protein translation.(49, 50) Deficiencies in protein synthesis may contribute to some cognitive deficits observed in Q331K mice,(44) including attenuated cue-induced freezing in the CCFC test in Q331K mice prior to surgery.(69) Although TBI is known to induce translational repression, upregulated expression of select proteins is thought to be important for mitigating the effects of trauma.(104) Our data support a model whereby Q331K mice are unable to mount a physiological response to TBI due to multiple deficits within the proteostasis network that likely include deficits in protein synthesis and folding (**Fig. 7**).

In addition to TBI,(13, 37–41) TDP-43 has been linked to other stress-related pathways in vitro(2–6, 11, 12) and in vivo.(15, 78). That TDP-43 undergoes a change in subcellular localization in response to a broad range of stressors suggests TDP-43 may serve as a general stress-sensor, a notion supported by our observation that TDP-43 re-distributes to the cytoplasm under conditions of sham surgery. Stress-induced subcellular re-distribution of TDP-43 is modulated by TDP-43 RNA-binding capacity and TDP-43 post-translational modification status, and this redistribution can coincide with mis-splicing signatures corresponding to TDP-43 loss of function.(3, 4, 7–9, 105) Here, we observe robust TBI-induced changes in subcellular TDP-43 localization in both WT and Q331K mouse brain tissue, including a ring-like TDP-43 staining pattern that has not been widely reported. Changes in TDP-43 localization were most prominent in the hypothalamus and the amygdala, the latter region having been linked to behavioral dysfunction in human sporadic ALS.(106) *A priori*, we expected TBI-induced changes in TDP-43 localization to coincide with splicing defects associated with TDP-43 loss-of-function.(4, 9, 82, 105) However, the splicing patterns of known TDP-43 targets were primarily influenced by TDP-43 mutation and largely unaffected by TBI. It is possible that emerging reporters of TDP-43-mediated cryptic splicing could be more sensitive for detection of TDP-43 loss-of-function in cells with altered TDP-43 localization in our TBI model.(107)

The outcomes of our study lend support to the hypothesis that ALS pathogenesis occurs through a multistep process, where steps may represent genetic, environmental and/or behavioral risk factors.(34, 35) Notably, we detected a synergistic effect of TDP-43 mutation and TBI on gene expression profiles that are relevant to ALS pathogenesis. Indeed, there is substantial overlap in GSEA hallmark gene sets in CNS tissues between Q331K TBI mice 24h post-TBI and human post-mortem ALS/FTLD-TDP cases,(45, 46) even though the Q331K TBI mice eventually recover, and the human cases represent end-stage disease. Our interpretation is that expression of mutant TDP-43 causes mice to become vulnerable to CNS challenge, resulting in dysregulation of key biological pathways that are also dysregulated during ALS/FTD pathogenesis. In Q331K mice, TBI-induced phenotypes that were observed 24h post-TBI, such as exaggerated neurological deficits (i.e., higher NSS), proteostasis-related gene upregulation, changes in *Ube3c* splicing and subcellular TDP-43 re-distribution, were largely restored to baseline levels 5-7 months post-TBI, consistent with this being a mild model of TBI.(43). Further, compensatory mechanisms, such as an upregulation of proteostasis-related genes, appear to be activated in Q331K mice (**Fig. 7**). Despite these protective measures, Q331K mice exhibited worse neurological outcomes in the acute phase post-TBI, indicating that stress-related pathways were vulnerable to dysfunctional TDP-43. This notion is further supported by a recent study demonstrating reduced stress granule formation in mutant TDP-43 mice exposed to hyperthermia.(15) We posit that repeated incidence of TBI and/or exposure to additional risk factors would lead to pathological and chronic degenerative phenotypes that persist months after the injury occurs (**Fig. 7**). Mathematical modeling of incidence data for human ALS cases with TDP-43 mutation indicated that disease pathogenesis may require up to five steps, where only one step is attributed to TDP-43 mutation.(34, 35) In the case of sporadic ALS, six steps are predicted for disease pathogenesis. Accordingly, environmental and behavioral risk factors likely play a significant role in ALS pathogenesis.(32, 33) While the TDP-43 Q331K mutation represents one step in the current study, TDP-43 dysfunction can also manifest in the absence of genetic mutations. For example, altered post-translational and transcriptional forms of TDP-43 that have been detected in sporadic ALS cases have also been linked to TDP-43 dysfunction.(108, 109)

### Conclusions

Outcomes of this study demonstrate that expression of dysfunctional TDP-43 predisposes mice to worse neurological outcomes following brain trauma. Deficiencies in multiple branches of the proteostasis network likely contribute to this vulnerability of mutant TDP-43 mice to TBI (**Fig. 8**). Therefore, targeting the proteostasis network should be a viable therapeutic direction for treating TDP-43 proteinopathies. For example, proteostasis could be enhanced by mitigating protein misfolding and stress through activation of heat-shock proteins and chaperones.(110) TDP-43 dysfunction can manifest through both genetic and non-genetic causes, where mutant TDP-43 expression altered the physiological response to TBI herein. We predict that TDP-43 dysfunction will similarly affect cellular response and recovery pathways associated with other forms of CNS challenge.(15) Further, repeated and/or additional incidences of CNS challenge are expected to increase the likelihood of developing a TDP-43 proteinopathy such as ALS (**Fig. 8**).

## Supporting information

SuppTableI

SuppTableII

SuppTableIII

SuppTableIV

SuppTableV

SuppTableVI

SuppTableVII

## List of abbreviations

Abbreviation: Description
AD: Alzheimer’s disease
ALS: amyotrophic lateral sclerosis
BCA: bicinchoninic acid
BSA: bovine serum albumin
Calr: calreticulin
CCFC: cued and contextual fear conditioning
CR: conserved region
CTE: chronic traumatic encephalopathy
Dnajc5: DnaJ heat shock protein family (Hsp40) member 5
ER: endoplasmic reticulum
ES: enrichment score
FOV: field of view
FTD: frontotemporal dementia
FTLD: frontotemporal lobar degeneration
Gid4: glucose-induced degradation protein 4 homolog
GO: gene ontology
GOBP: Gene Ontology Biological Process
GSEA: Gene Set Enrichment Analysis
GT: genotype
HECT: homologous to the E6-AP Carboxyl Terminus
Hspa4: heat shock 70 kDa protein 4
IACUC: Institutional Animal Care and Use Committee
IDR: intrinsically disordered region
INJ: Injury (i.e. sham or TBI)
Kcnip2: potassium voltage-gated channel interacting protein 2
LATE: limbic-predominant age-related TDP-43 encephalopathy
LC-MS/MS: liquid chromatography tandem mass spectrometry
Manf: mesencephalic astrocyte-derived neurotrophic factor
MCL: Markov Clustering
MSigDB: molecular signatures database
MTOR1: mammalian target of rapamycin complex 1
N/C: nuclear-to-cytoplasmic
NES: normalized enrichment score
NMDA: N-methyl-D-aspartate
NSS: neurological severity score
Pacrgl: parkin coregulated like
PCA: principal component analysis
Pdia3: protein disulfide isomerase family A, member 3
Pdia6: protein disulfide isomerase family A, member 6
PFA: paraformaldehyde
PISA: proteome integral solubility alteration
Ppp3ca: protein phosphatase 3 catalytic subunit alpha
pre-mRNA: pre-messenger RNA
PSM: peptide-spectrum match
PVDF: polyvinylidene fluoride
RNA-seq: bulk RNA-sequencing
RNP: ribonucleoprotein
Rplp1: large ribosomal subunit protein P1
Rps27a: ribosomal protein S27a
RT-PCR: Reverse transcription polymerase chain reaction
Slc6a6: solute carrier family 6 member 6
Sort1: Sortilin-1
Spop: speckle-type POZ protein
Ssr1: signal sequence receptor subunit 1
TAMC: Transgenic Animal Modeling Core
TBI: traumatic brain injury
TCP-1: cytosolic T-complex protein 1
TDP-43: TAR DNA-binding protein 43
TMT: tandem mass tag
TRiC/CCT: eukaryotic TCP-1 ring complex or chaperonin containing TCP-1
Tsn: translin
Uba52: ubiquitin A-52
Ube3c: ubiquitin protein ligase E3C
UPLC: UltraPerformance LC
WT: wild-type
Xbp1: active X-box binding protein 1
Xbp1s: spliced active X-box binding protein 1
Xbp1u: unspliced active X-box binding protein 1

## Declarations

### Ethics approval and consent to participate

Not applicable.

### Consent for publication

Not applicable.

### Availability of data and materials

Mass spectrometry proteomics data are available via ProteomeXchange with identifier PXD068825 and 10.6019/PXD068825.Submission of RNAseq dataset to GEO are in progress at the time of this manuscript submission.

### Authors’ contributions

**MSR** performed all the mass spectrometry experiments and analyzed the resulting data, performed most of the RNAseq analyses and was a major contributor in writing and preparing the manuscript; **MFM** performed the TDP-43 localization analyses, contributed to the RNA-related analyses, and created multiple figures in the manuscript; **JZ** performed animal surgeries, performed the CCFC analyses, and acquired the confocal microscopy images; **KO** performed the RT-PCR analyses; **EAW** isolated RNA from mouse tissue and coordinated the RNA sequencing and provided some assistance in the analysis; **DC** maintained the mouse colony and assisted in the animal studies; **KS** assisted in the animal studies and performed immunofluorescence; **JB** performed the NSS tests and helped with surgeries; **HM** performed the AlphaFold analyses and contributed to the RT-PCR analyses; **MNA** performed the polysome profiling studies together with MSR; **FM** performed the PyMOL analyses and interpreted the results; **ART** and **SM** assisted in the CCFC study; **JAN** assisted with confocal microscopy; **NH** contributed to the experimental design for the TBI surgeries and analyzed the animal behavior data; **ODK** performed the tests of differential gene expression for the RNAseq data and was a major contributor to the omics analyses. **DAB** conceived and directed the project, analyzed data and wrote the manuscript with contributions from co-authors.

## Acknowledgements

We are grateful for services provided by UMass Chan core facilities including the Transgenic Animal Modeling Core, Mouse Behavioral Core, Mass Spectrometry Facility and Molecular Biology Core Labs. We thank Dr. Joel Richter for use of equipment and advice related to polysome profiling; Drs. Patrick Sullivan, Hemendra Vekaria, Myriam Heiman, Jack Humphrey, Sami Barmada, Christine Vande Velde and Clotilde Lagier-Tourenne and members of their labs for valuable advice; and current and former members of Bosco lab, especially Dr. Ozge Yildiz and Jonathan Jung for help on this project.

## Authors’ information

Not applicable.

## Additional file 1. Supplemental Figures (Fig. S1 thru S11)

**Figure S1.**
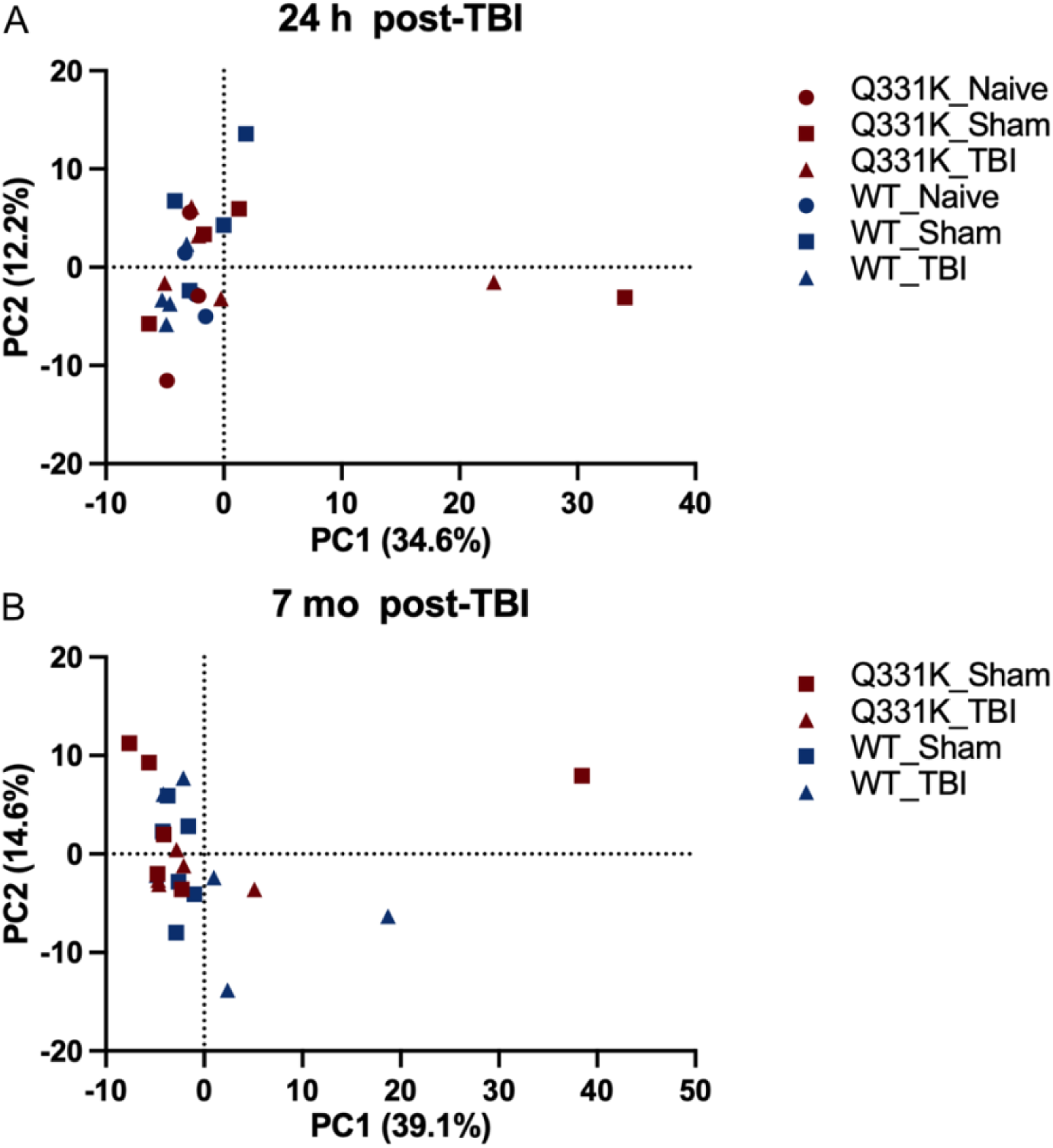
Principal component analysis (PCA) identifies outliers in RNAseq datasets. PCA analysis using 1000 genes with the highest variance for the 24 h post-TBI **(A)** and 7 mo post-TBI (**B**) datasets. Genes were selected based on their variance (sigma^2^) after adjusting for the mean-variance trend by fitting the expression data with the limma functions voom and lmFit, both using a design with a single group.

**Figure S2.**
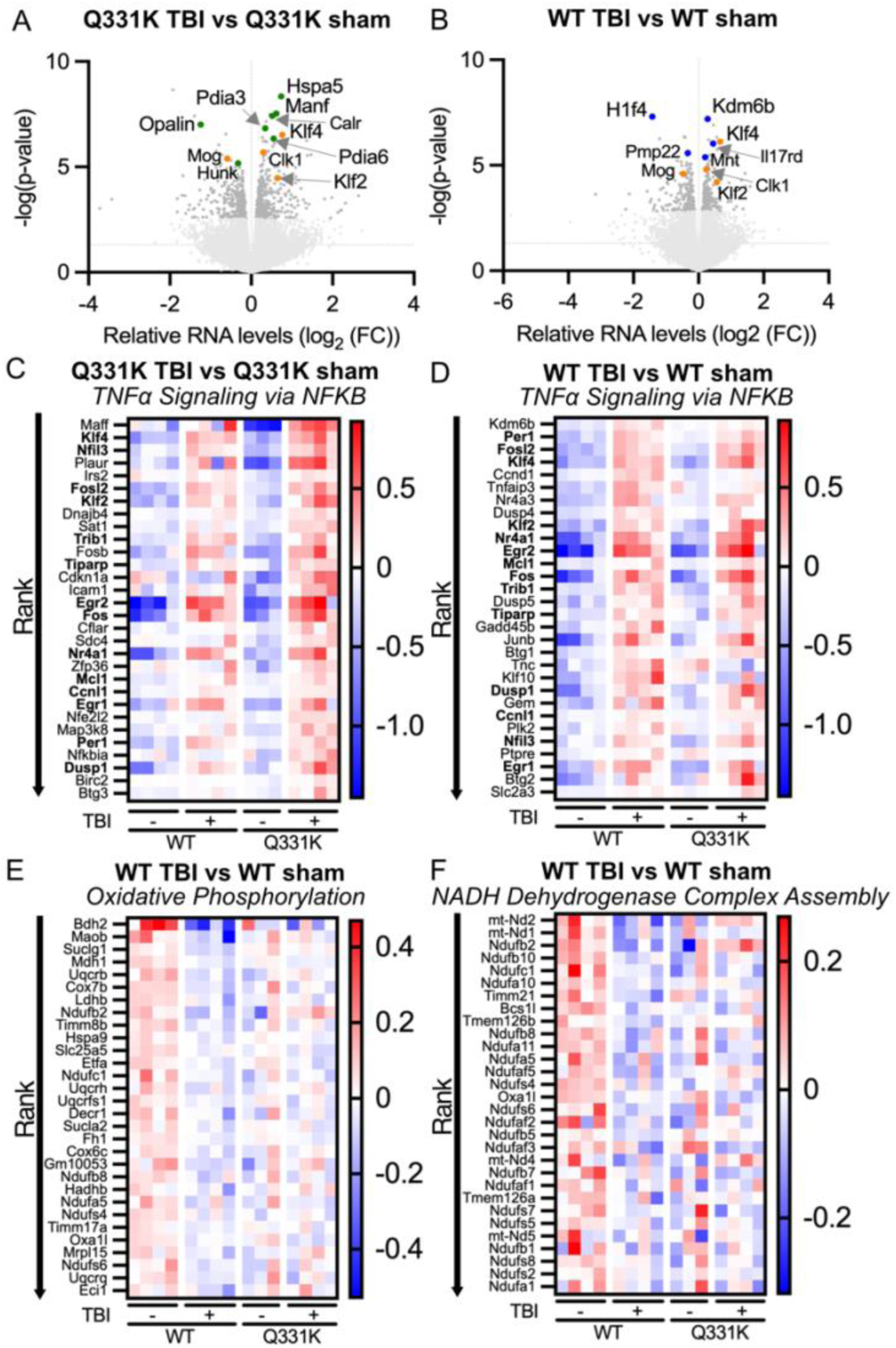
Overlapping and differential gene expression profiles in WT versus Q331K mice after TBI. **(A)** Volcano plot of transcriptional changes from RNAseq analyses for Q331K mice under conditions of TBI versus sham surgery. **(B)** Volcano plot of transcriptional changes from RNAseq analyses for WT mice under conditions of TBI versus sham surgery. **(A,B)** Hits with FDR <0.1 are shown in red. Orange dots highlight top gene hits (*Klf2*, *Klf4*, *Mog*, *Clk1*) resulting from TBI, irrespective of genotype. Blue dots denote top gene hits specifically in the WT TBI condition whereas green dots are top gene expression changes specifically in the Q331K TBI condition. **(C,D)** Heatmap showing the top 30 ranked genes in the top up-regulated gene set for hallmark “TNFalpha signaling via NFKb” for Q331K TBI versus Q331K sham (C) and WT TBI versus WT sham (D). Fourteen genes (in bold) are significantly altered under “TNFalpha signaling via NFKb” for both the Q331K TBI and WT TBI conditions relative to their respective sham group. **(E,F)** Heatmap showing the top 30 ranked genes in the top down-regulated gene set in hallmark collection, “Oxidative Phosphorylation” (E), and Gene Ontology Biological Process collection, “NADH Dehydrogenase Complex Assembly” (F), for the WT TBI versus WT sham comparison. The log2(fold change) relative to the mean expression is shown for each sample.

**Figure S3.**
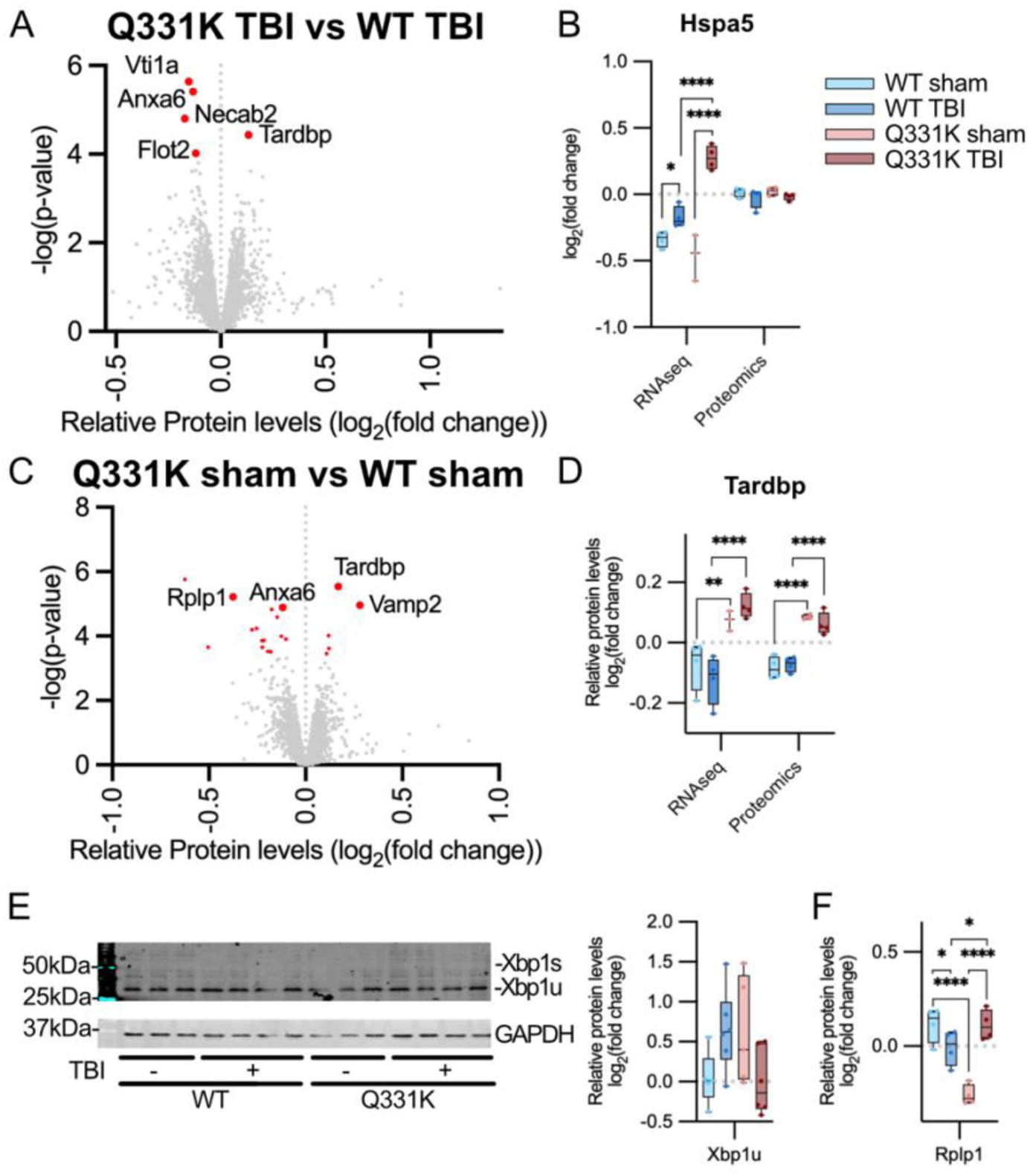
Proteomic analyses identify protein changes altered by TBI and mutant TDP-43 genotype. **(A)** Volcano plot showing relative protein level changes between Q331K TBI and WT TBI cohorts. Hits with FDR <0.1 are highlighted in red. **(B)** Hspa5 is unchanged at the protein level by quantitative proteomics. **(C)** Volcano plot showing relative protein level changes between Q331K sham and WT sham cohorts. Hits with FDR <0.1 are highlighted in red. **(D)** Tardbp is significantly elevated in Q331K mouse brain by transcriptomics and proteomics analyses. **(E)** Total levels of Xbp1 are unchanged and there is no detection of spliced Xbp1 by Western analysis. The expected molecular weight of spliced Xbp1 (Xbp1s) is 54 kDa and of unspliced Xbp1 (Xbp1u) is 29 kDa. Note, the band located at ∼35 kDa is nonspecific. **(F)** Protein levels of Rplp1 as determined by quantitative proteomics. Nominal p-value = * <0.05; **<0.01, ****<0.0001.

**Figure S4.**
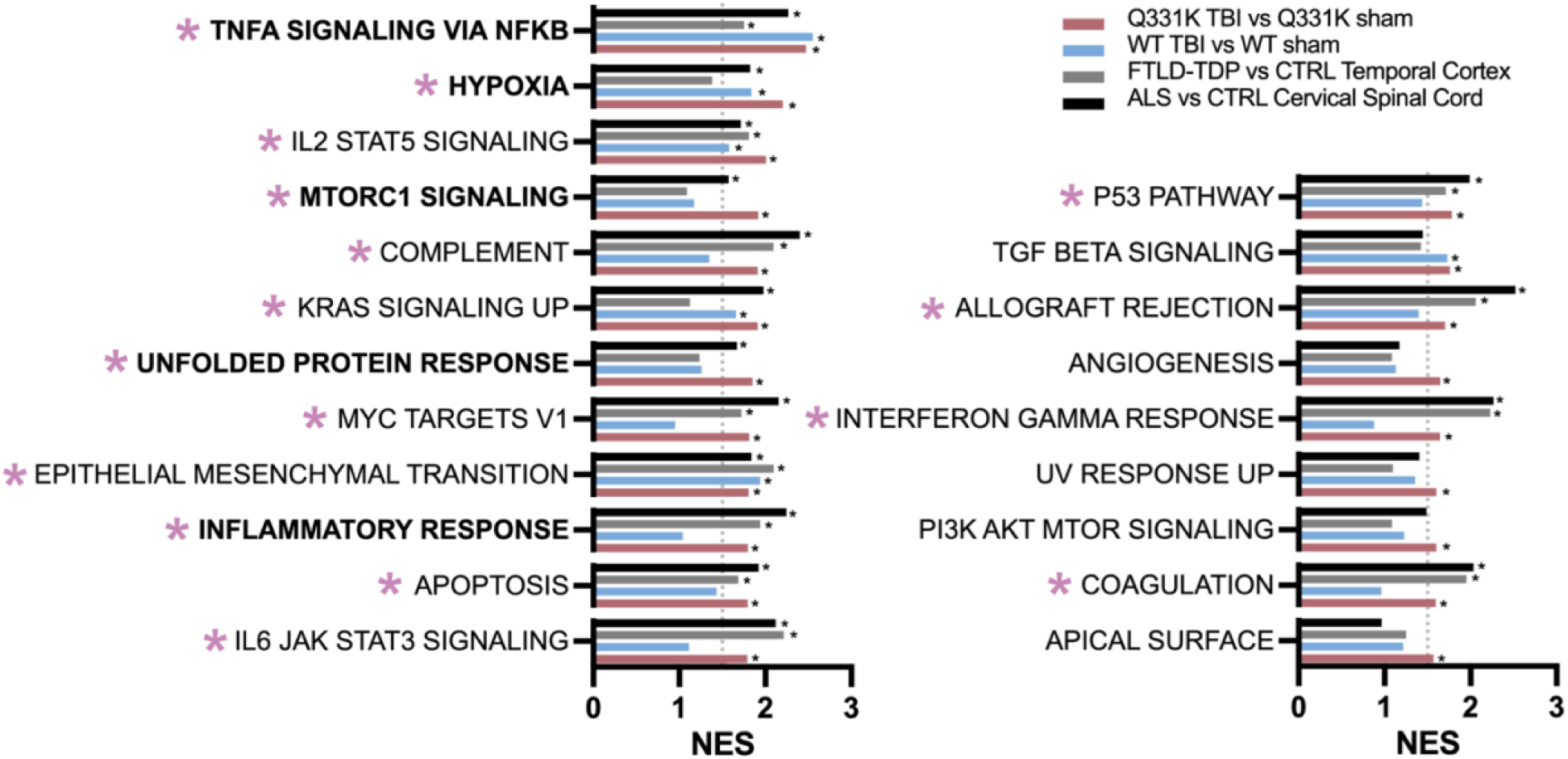
Comparison of published human ALS and FTLD post-mortem datasets with current TBI mouse study. Directional p-values were calculated from published datasets reported in Humphrey et al,(46) and Hasan et al (45) and subjected to GSEA using the hallmark database as described in methods for RNAseq. Asterisks to the right of the bars denote FDR<0.05 for the indicated comparison defined in the key. The dotted line indicates NES of 1.5. Gene sets with a pink asterisk highlight those that are significant for the Q331K TBI versus Q331K sham comparison (pink bars), and at least one of the human datasets (grey and black bars). There are 16/21 gene sets that meet this criterion, and only 5/21 gene sets that are significant for the WT TBI versus WT sham comparison (blue bars) and at least one of the human datasets. Gene set names in bold are discussed in the main text.

**Figure S5.**
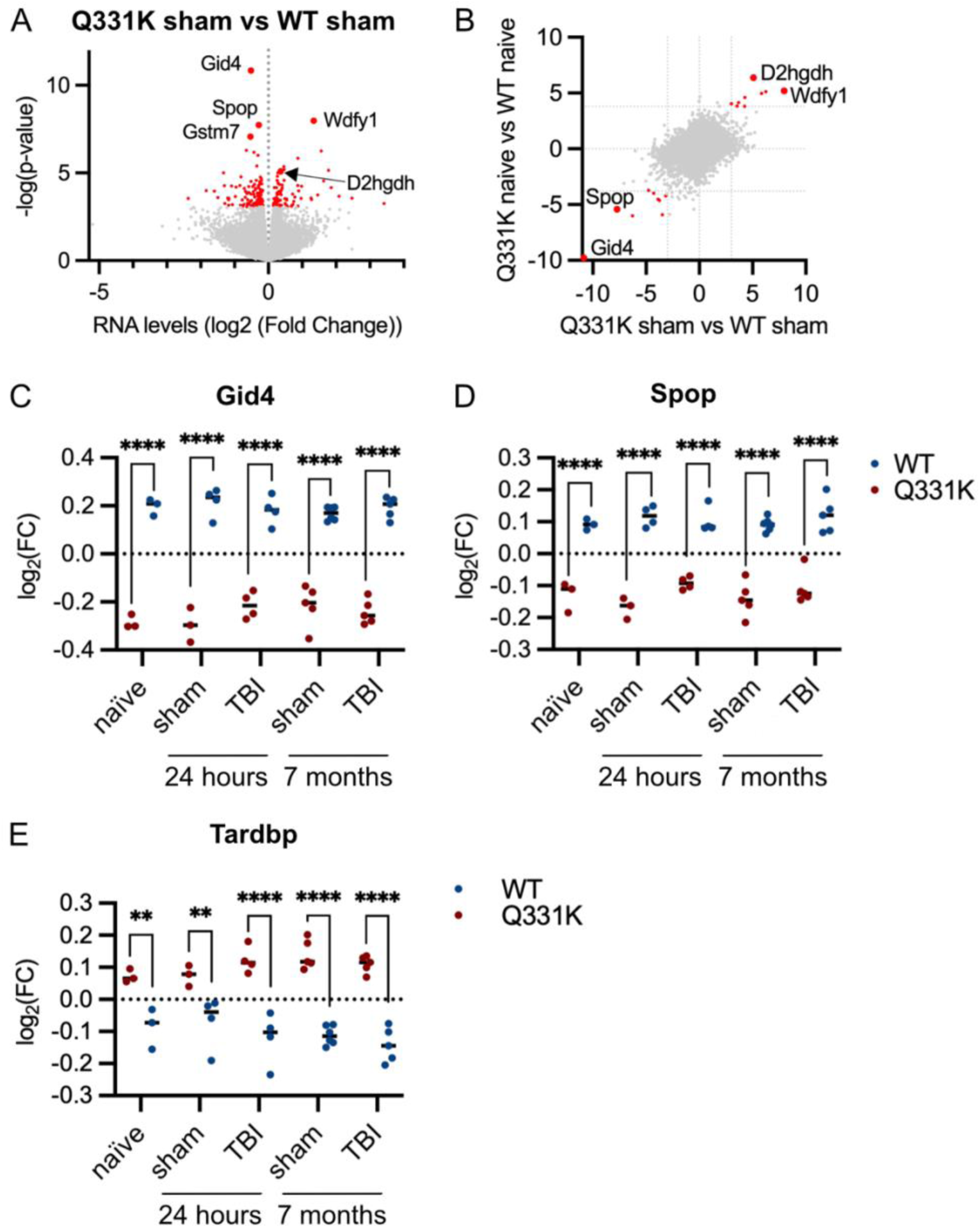
Omics analyses identify Q331K-specific changes to the transcriptome. **(A)** Volcano plot with the significant up- and down-regulated transcripts in red for the Q331K sham versus WT sham comparison. **(B)** Directional p-value plot overlaying the Q331K sham versus WT sham datasets with the Q331K naive versus WT naive datasets to identify reproducible mutant TDP-43-specific changes, which are highlighted in red. Directional p-value = -log_10_(p-value)*sign of log_2_(fold change) **(C-D)** Plots of the most significant Q331K-specific down-regulated genes, *Spop* and *Gid4*, which are both involved in the ubiquitin-proteosome pathway. **(E)** Elevated *Tardbp* is observed in all Q331K vs WT comparisons. For C-E, time post-surgery is indicated below. Nominal p-value = **<0.01, ****<0.0001.

**Figure S6.**
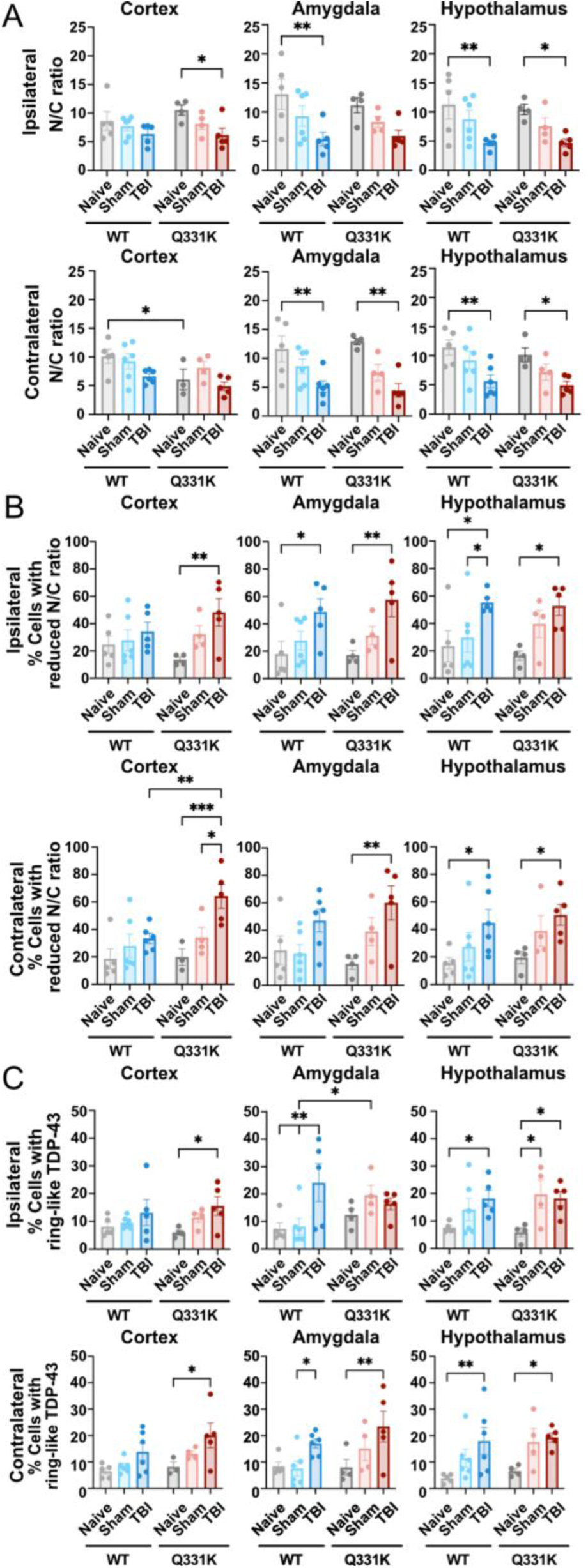
Changes in TDP-43 subcellular localization in mouse brain tissue 24h post-surgery. **(A)** Nuclear-to-cytoplasmic (N/C) ratio of TDP-43 signal in naïve mice or 24h post-surgery (sham or TBI) mice determined from immunofluorescence analysis of tissue sections from the cortex (left), amygdala (middle), and hypothalamus (right), either ipsilateral (top) or contralateral (bottom) to the TBI site. **(B)** Percent of cells that exhibit reduced TDP-43 N/C ratios in the tissues from (A). **(C)** Percent of cells that exhibit “ring-like” TDP-43 distribution in the tissues from (A). Each dot represents data from an individual mouse, with n=4-6 mice per cohort. Statistics was performed using two-way ANOVA followed by Tukey’s multiple comparisons test between conditions. *p<0.05, **p<0.01, ***p<0.001, ****p<0.0001

**Figure S7.**
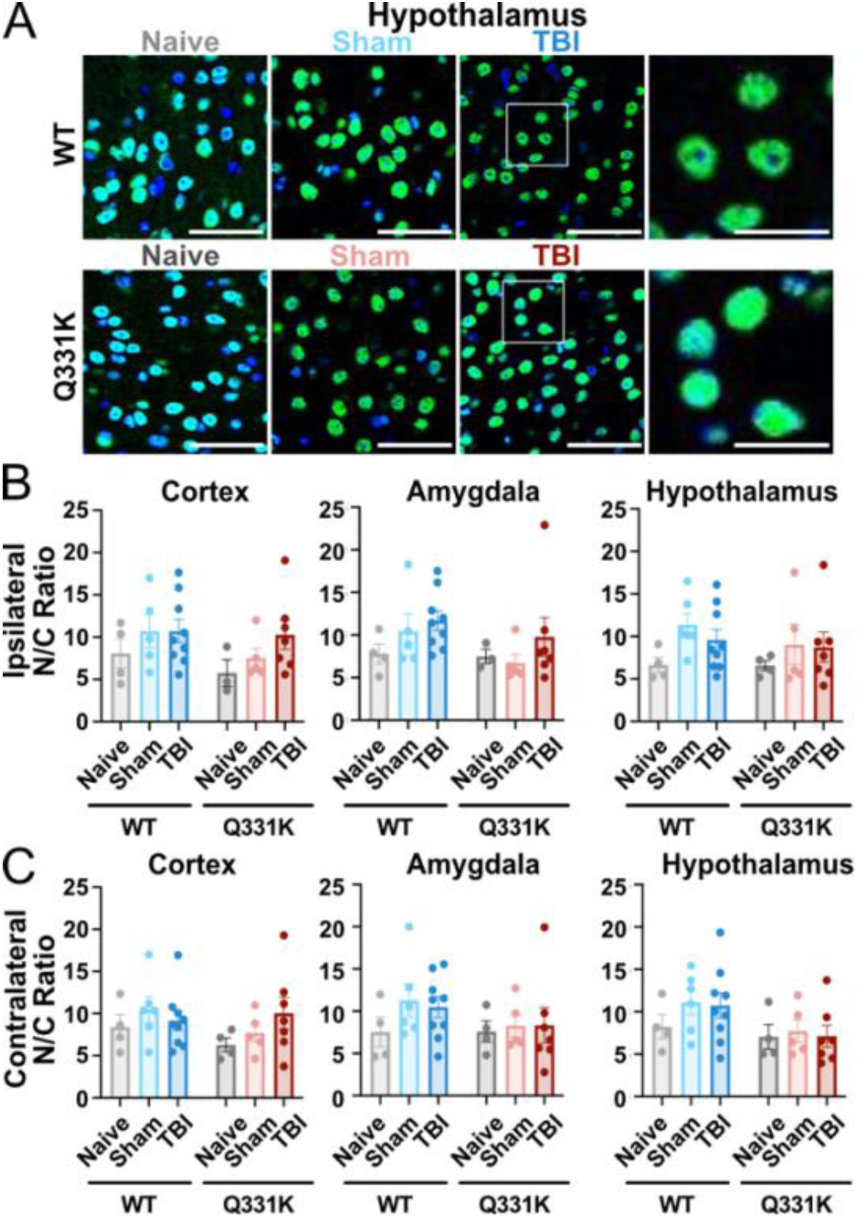
Lack of significant TDP-43 subcellular alteration in mouse brain tissue 5 months post-surgery. **(A)** Confocal immunofluorescence images of hypothalamus brain sections from WT and Q331K mice, either without surgery (naïve) or 5-months post-surgery (sham or TBI). Sections are stained for TDP-43 (green) and counterstained with DAPI (blue). Scale bars=20 um. **(B,C)** Quantification of the nuclear-to-cytoplasmic (N/C) ratio of TDP-43 signal in WT and Q331K mice 5-months post-surgery in the cortex, amygdala and hypothalamus, either ipsilateral (B) or contralateral with respect to the impact site. Each dot represents data from an individual mouse, with n=4-9 mice per cohort. Statistics was performed using two-way ANOVA followed by Tukey’s multiple comparisons test between conditions.

**Figure S8.**
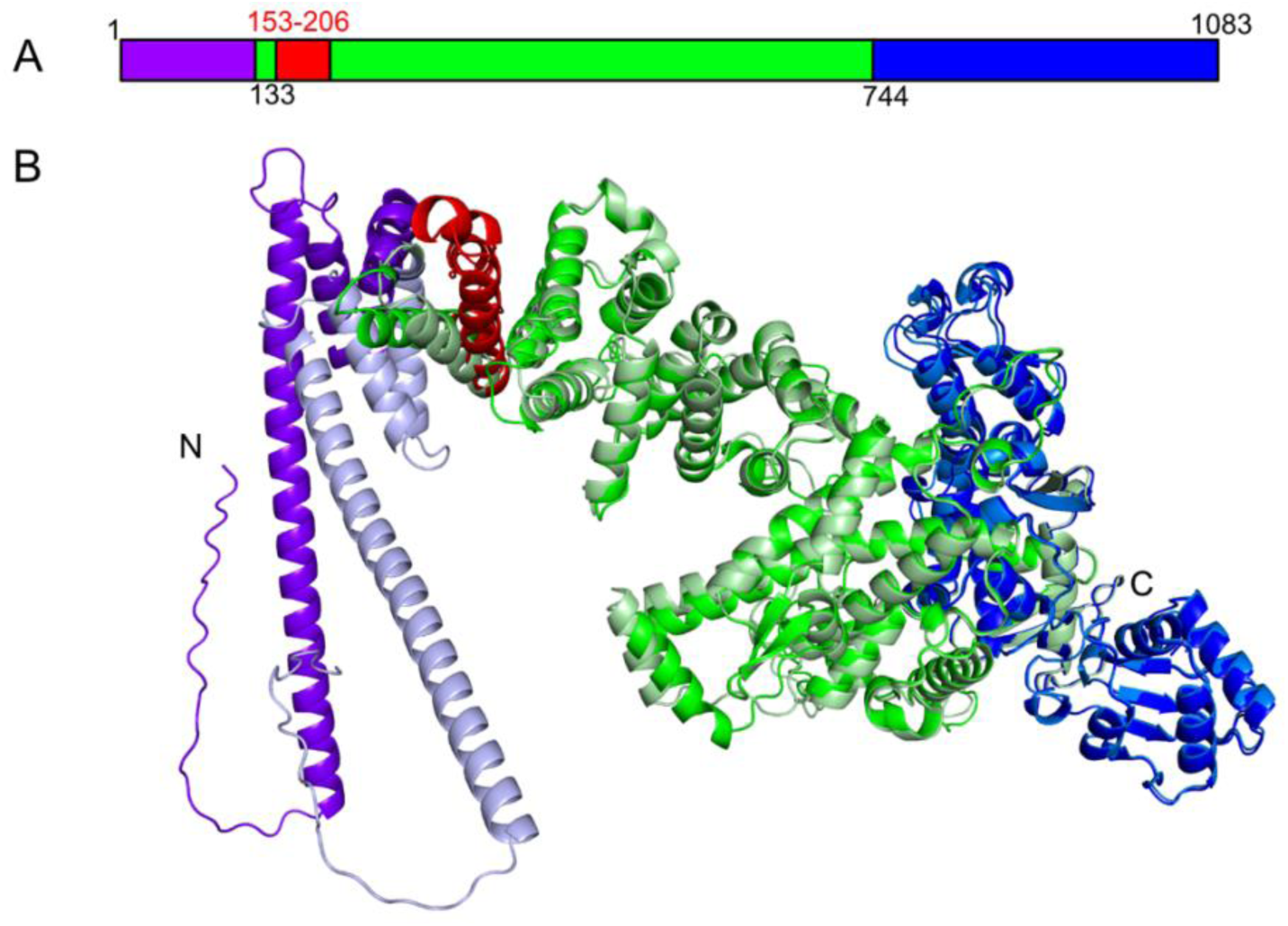
Structural analysis of TDP-43-relevant splice variants using AlphaFold and PyMOL. **(A)** Schematic representation of the ubiquitin-protein ligase E3C (Ube3c) protein highlighting distinct domains. Residue numbers are indicated at the start of each domain. The ubiquitin-binding domain (residues 1–132) is shown in purple, the second domain (residues 133–743) in green, and the catalytic Homologous to the E6-AP Carboxyl Terminus (HECT) domain (residues 744–1083) in blue. Residues corresponding to exon 6 (residues 153–206) are highlighted in red. **(B)** Overlay of AlphaFold-predicted structures for Ube3c containing exon 6 (domains in purple, green, and blue) with the predicted structures for Ube3c lacking exon 6 (domains in light purple, light green, and light blue). Structural alignments were performed in PyMOL, yielding an RMSD of 2.3 Å for alignment of only the ubiquitin-binding domain and an RMSD of 8.5 Å for the entire protein.

**Figure S9.**
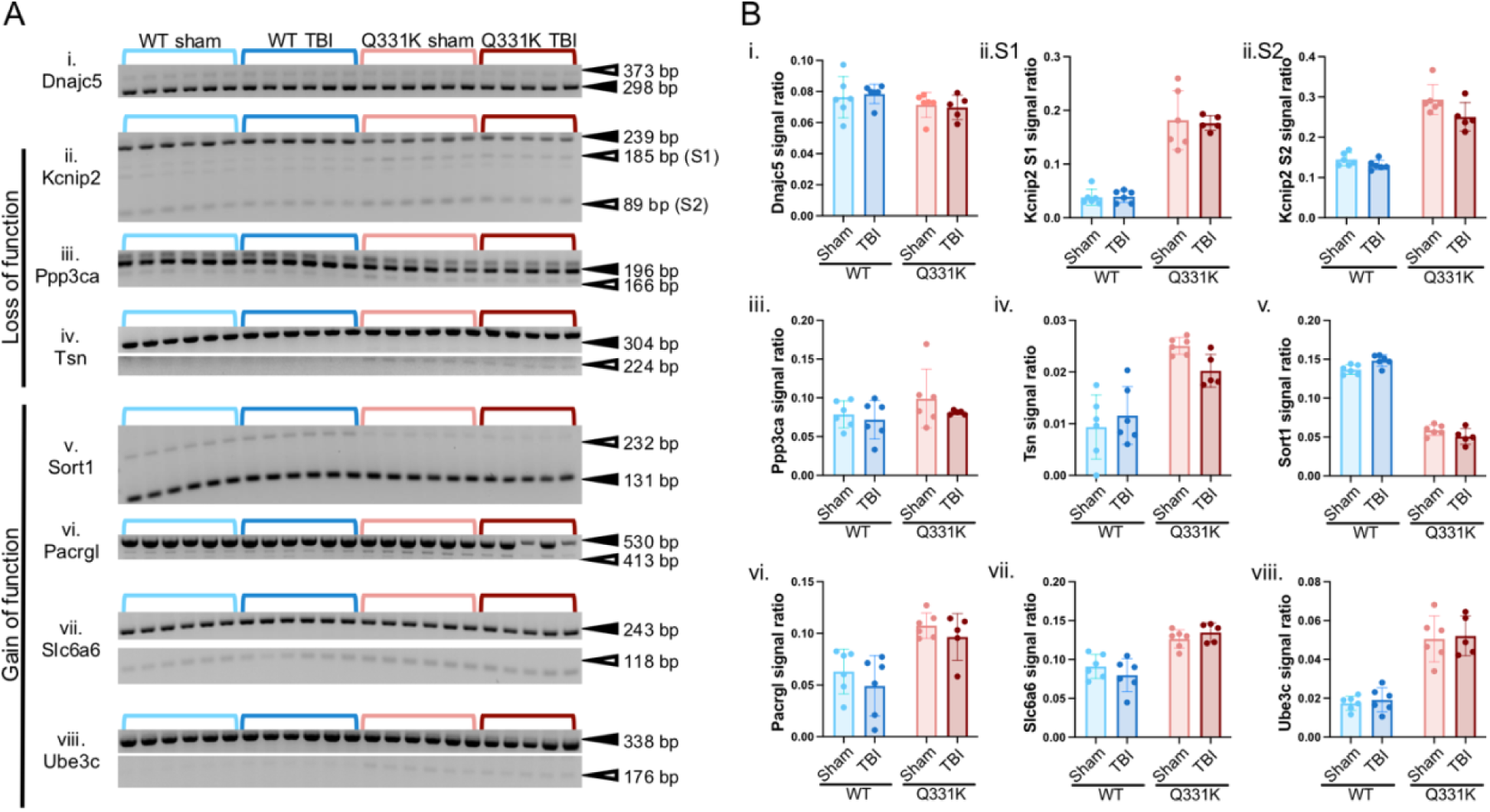
Alternative splicing targets of TDP-43 in WT and Q331K mice at 7 months post-injury. **(A)** Splice variants of known TDP-43 RNA targets (i-vi) detected by RT-PCR analysis in WT and Q331K mice 24h after either TBI or sham surgery. The PCR primers, exons of interest and references for the primers used in this study are shown in **Supp Table VI**. Splicing patterns that are consistent with TDP-43 loss of function or gain of function are indicated. Common and mis-spliced variants are denoted by, respectively, closed and open arrowheads with the corresponding size (bp=base pairs) of the PCR product. **(B)** Densitometry analysis of the gels in (A) was used to determine the ratio of the mis-spliced/common PCR product for each gene of interest. Each dot in the graph represents a separate mouse.

**Figure S10.**
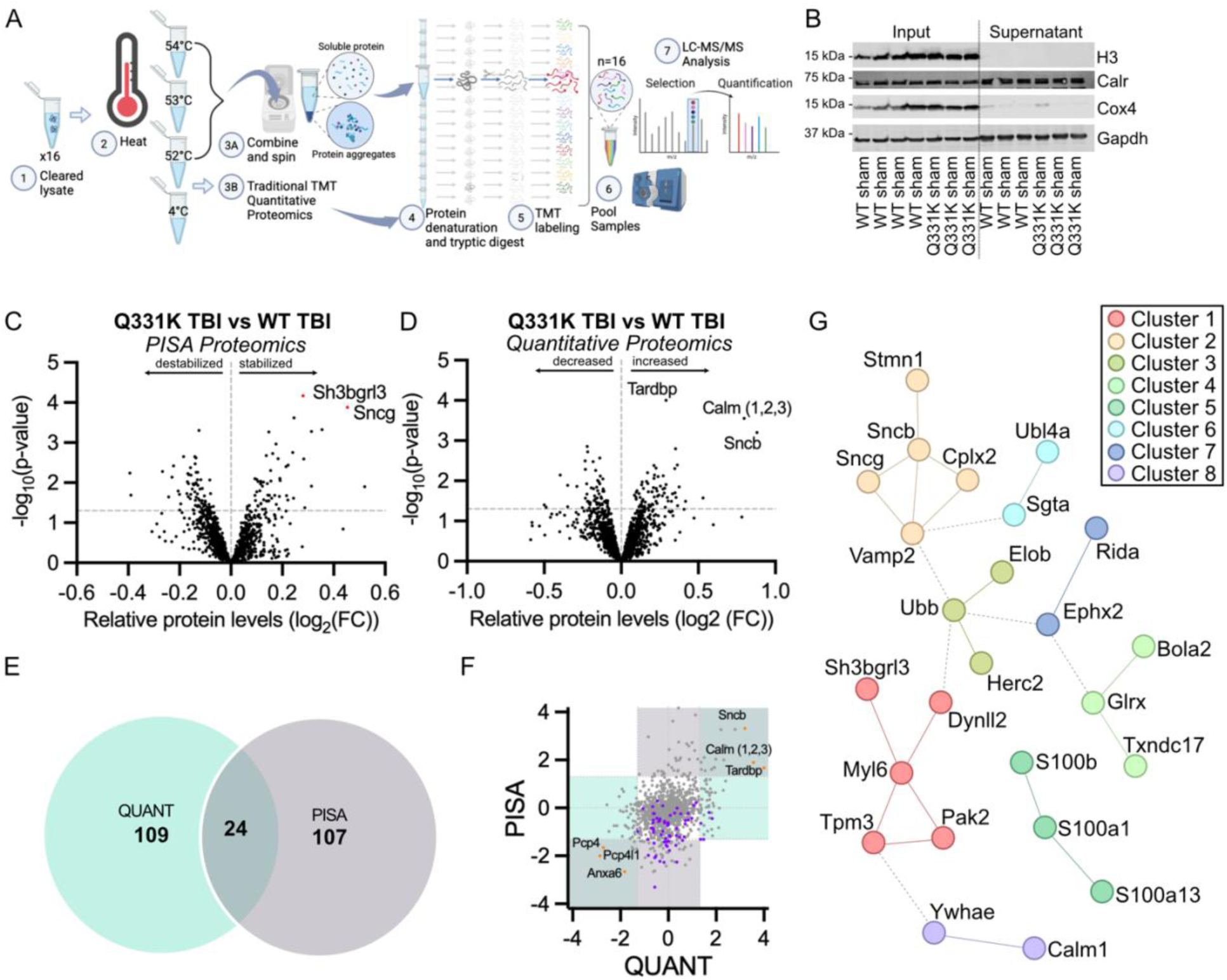
Quantitative proteomics of organelle-depleted cytoplasmic brain lysates. (**A**) PISA protocol for identifying proteins with altered thermostability between conditions: (1) Lysates are generated with mild, non-denaturing conditions and homogenization. Lysates are cleared by centrifugation. (2) Each sample is divided into four samples, where one is placed on ice for traditional quantitative proteomics and three are subjected to heat for protein denaturation. (3A) Aggregated protein is removed by centrifugation. (3B) The sample for traditional TMT quantitative proteomics remains on ice while samples for PISA are heated. (4) The soluble protein is digested with trypsin and labeled (5) with isobaric TMTpro-18plex tags. (6) Tagged samples are pooled, generating two sample sets (one from PISA and one from traditional quantitative proteomics) and subjected to mass spectrometry for quantification (7). Created with biorender.com. (**B**) Western analyses of organelle depleted supernatant (supernatant) compared to the whole brain sample input (input). (**C-D**) Volcano plot of protein changes identified by traditional quantitative proteomics (**C**) and PISA (**D**) for Q331K TBI versus WT TBI conditions. (**E**) Overlapping hits between quantitative proteomics and PISA are assessed to distinguish protein level changes from thermostability changes. (**F**) Directional p-value plot of Q331K TBI versus WT TBI for PISA and QUANT (quantitative proteomics). Purple dots indicate ribosomal proteins. Orange dots are highlighting top hits found in both PISA and QUANT. (**G**) Clustering using the Markov Cluster Algorithm or “Q331K All” versus “WT All” (defined in Figure 7 legend) for the stabilized proteins passing FDR adjustment (FDR <0.1). Only proteins identified as part of a cluster are shown. Cluster 1: Smooth Muscle Contraction; Cluster 2: Synaptic vesicle endocytosis, Synaptobrevin 2-SNAP-25-syntaxin-1a-complexin II complex; Cluster 3: Inactivation of Granulocyte colony-stimulating factor (G-CSF) signaling; Cluster 4: Protein-disulfide reductase nicotinamide adenine dinucleotide phosphate (NAD(P)) activity; Cluster 5: S-100/Intestinal calcium-binding protein (ICaBP) type calcium binding domain; Cluster 6: ER membrane insertion complex; Cluster 7: Epoxide hydrolase 2 (Ephx2), reactive intermediate imine deaminase A (Rida); Cluster 8: Negative regulation of calcium ion export across plasma membrane. Protein-protein interaction enrichment p-value = 2.94e^-06^.

**Figure S11.**
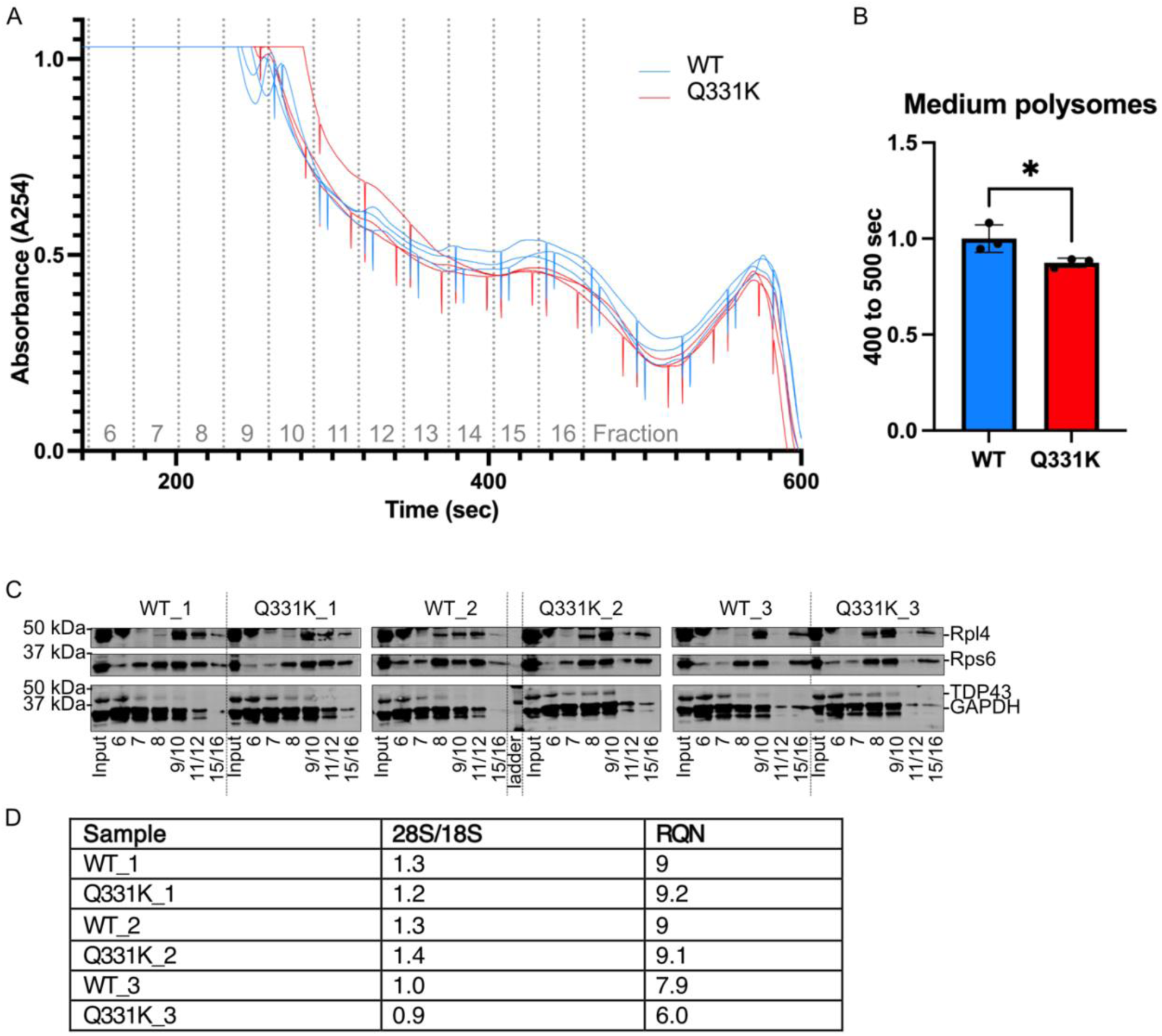
Polysome profiling of cytoplasmic fractions confirm the presence of intact ribosomes. **(A)** Graph of sucrose gradient fractions post-ultracentrifugation. The prominent peaks are at fractions 9/10, fractions 11/12, and fractions 15/16. **(B)** The area under the curve was calculated for all samples at 400 sec to 500 sec to estimate relative amount of medium polysomes, as this was the only ribosomal peak consistently observed in all samples. * = p-value <0.05. **(C)** Western analyses of the protein precipitated fractions of the polysome profile shown in A. Numbers indicate specific fractions analyzed. **(D)** Fragment analysis of RNA isolated from cytoplasmic fractions for each sample subjected to polysome profiling using the Agilent Fragment Analyzer.

**<u>Additional file 2. FIJI Macro for Per FOV N/C Analysis</u>**. Macro script to be used in FIJI/ImageJ to complete TDP-43 localization analysis on a per FOV level. DAPI signal is used to define and separate the nuclear and non-nuclear regions where TDP-43 fluorescent signal is measured to determine the N/C ratio.

**<u>Additional file 3. CellProfiler Pipeline for Per Cell N/C Ratio and Ring-like Analysis.</u>** CellProfiler Pipeline used for per cell analyses of TDP-43 fluorescent signal including N/C ratio and ring-like TDP-43 signal. DAPI signal is used to identify individual cells, as well as define nuclear and cytoplasmic regions for N/C ratio analyses. For ring-like analysis, the nuclear region is binned into 3 rings to report on distribution of TDP-43 signal.

**Additional file 4, Supp. Table I. Statistical output of RNAseq analyses**. For all comparisons, logFC refers to log2(fold change), PValue refers to nominal p-value, and FDR refers to adjusted p-value. Nominal p-values <0.05 are indicated in orange and FDR < 0.1 in red. (**IA**) Statistical analyses for the 4 mo old mouse cohorts including the 24 h post-surgery (TBI and sham) and naïve groups. (**IB**) Statistical analyses for the 7 mo post-surgery cohorts.

**Additional file 5, Supp.Table II. GSEA analyses for all comparisons from the RNAseq analysis**. All gene set hits that passed the cutoff criteria (NES > 1.5, FDR > 0.05) are listed. Each tab is a unique group comparison. In each tab, the top upregulated gene sets are beneath the purple-shaded header on the left, while the top downregulated gene sets are beneath the blue-shaded header on the right. Results of hallmark and GOBP are shown. (**IIA**) List of abbreviations used in the file. (**IIB-E**) Gene set hits for the 24h-post surgery (TBI and sham). (**IIF**) Gene set hits for Q331K naïve mice relative to WT naïve mice at 4 months of age. (**IIG-IIJ**) Detailed gene lists for the top pathway hits for the 24h group comparisons. (**IIK-IIN**) Gene set hits for each group comparison at 7 mo post-surgery.

**Additional file 6, Supp. Table III. Statistical analyses of whole brain quantitative proteomics from Q331K and WT mice with surgery**. Unadjusted p-values ≤ 0.05 are highlighted red (columns H-K). Adjusted p-values ≤ 0.1 are highlighted in red (columns L-O). Coef. refers to Log_2_(fold change); P.value indicates nominal p-value; P.value.adj. represents adjusted p-value (FDR).

**Additional file 7, Supp. Table IV. Overlap of published omics datasets with the current study.** Includes all identified proteome and transcriptome level changes in Q331K relative to WT mice from the current study. Columns B-D refer to datasets in the present study. RNAseq_Sham refers to the Q331K sham versus WT sham comparison. RNAseq_Naive refers to Q331K naïve versus WT naïve at 4 months of age. Proteomics_Sham refers to Q331K sham versus WT sham whole brain proteomics. Columns E/F refer to results published by White et al.(44).

**Additional file 8, Supp. Table V. RT-PCR primers.** Sequence information for primers that were used to identify alternative splicing for targets of TDP-43 in **Figures 5** and **S8**.

**Additional file 9, Supp. Table VI. Significant outcomes from quantitative proteomics on organelle-depleted cytoplasmic brain lysates.** (**VIIA**) PISA quantitative proteomics results. (**VIIB**) Traditional quantitative proteomics results (i.e. not exposed to heat). Unadjusted p-values ≤ 0.05 are highlighted red (columns K-Q). Adjusted p-values ≤ 0.1 are highlighted in red (columns R-X). Coef. refers to Log_2_(fold change); P.value indicates nominal p-value; P.value.adj. represents adjusted p-value (FDR).

**Additional file 10, Supp. Table VII. GSEA analyses of quantitative proteomics outcomes for organelle-depleted cytoplasmic brain lysates.** All gene set hits that passed the cutoff criteria (NES > 1.5, FDR > 0.05) are listed. Each tab is a unique group comparison. In each tab, the top upregulated gene sets are beneath the purple-shaded header on the left, while the top downregulated gene sets are beneath the blue-shaded header on the right. Results of hallmark and GOBP are shown. (**VIIIA-D**) Gene set hits for samples processed using PISA. (**VIIIE-H**) Gene set hits from the traditional quantitative proteomics analyses.

## References

1. Buratti E, Baralle FE. Multiple roles of TDP-43 in gene expression, splicing regulation, and human disease. Front Biosci. 2008;13:867–78.

2. Dewey CM, Cenik B, Sephton CF, Dries DR, Mayer P, 3rd, Good SK, et al. TDP-43 is directed to stress granules by sorbitol, a novel physiological osmotic and oxidative stressor. Molecular and cellular biology. 2010.

3. Duan L, Zaepfel BL, Aksenova V, Dasso M, Rothstein JD, Kalab P, et al. Nuclear RNA binding regulates TDP-43 nuclear localization and passive nuclear export. Cell Rep. 2022;40(3):111106.

4. Klim JR, Williams LA, Limone F, Guerra San Juan I, Davis-Dusenbery BN, Mordes DA, et al. ALS-implicated protein TDP-43 sustains levels of STMN2, a mediator of motor neuron growth and repair. Nat Neurosci. 2019;22(2):167–79.

5. Tischbein M, Baron DM, Lin YC, Gall KV, Landers JE, Fallini C, et al. The RNA-binding protein FUS/TLS undergoes calcium-mediated nuclear egress during excitotoxic stress and is required for GRIA2 mRNA processing. J Biol Chem. 2019;294(26):10194–210.

6. Weskamp K, Tank EM, Miguez R, McBride JP, Gomez NB, White M, et al. Shortened TDP43 isoforms upregulated by neuronal hyperactivity drive TDP43 pathology in ALS. J Clin Invest. 2020;130(3):1139–55.

7. Perez-Berlanga M, Wiersma VI, Zbinden A, De Vos L, Wagner U, Foglieni C, et al. Loss of TDP-43 oligomerization or RNA binding elicits distinct aggregation patterns. EMBO J. 2023;42(17):e111719.

8. Zhang S, Vazquez-Sanchez S, Lu S, Baughn MW, Lim J, Oung S, et al. Acetylation of lysine 82 initiates TDP-43 nuclear loss of function by disrupting its nuclear import. bioRxiv. 2024:2024.09.04.611121.

9. Necarsulmer JC, Simon JM, Evangelista BA, Chen Y, Tian X, Nafees S, et al. RNA-binding deficient TDP-43 drives cognitive decline in a mouse model of TDP-43 proteinopathy. Elife. 2023;12.

10. Dos Passos PM, Hemamali EH, Mamede LD, Hayes LR, Ayala YM. RNA-mediated ribonucleoprotein assembly controls TDP-43 nuclear retention. PLoS Biol. 2024;22(2):e3002527.

11. Liu-Yesucevitz L, Bilgutay A, Zhang YJ, Vanderwyde T, Citro A, Mehta T, et al. Tar DNA binding protein-43 (TDP-43) associates with stress granules: analysis of cultured cells and pathological brain tissue. PloS one. 2010;5(10):e13250.

12. McDonald KK, Aulas A, Destroismaisons L, Pickles S, Beleac E, Camu W, et al. TAR DNA-binding protein 43 (TDP-43) regulates stress granule dynamics via differential regulation of G3BP and TIA-1. Human molecular genetics. 2011.

13. Anderson EN, Gochenaur L, Singh A, Grant R, Patel K, Watkins S, et al. Traumatic injury induces Stress Granule Formation and enhances Motor Dysfunctions in ALS/FTD Models. Human molecular genetics. 2018.

14. Alami NH, Smith RB, Carrasco MA, Williams LA, Winborn CS, Han SSW, et al. Axonal transport of TDP-43 mRNA granules is impaired by ALS-causing mutations. Neuron. 2014;81(3):536–43.

15. Dubinski A, Gagne M, Peyrard S, Gordon D, Talbot K, Vande Velde C. Stress granule assembly in vivo is deficient in the CNS of mutant TDP-43 ALS mice. Human molecular genetics. 2023;32(2):319–32.

16. Gopal PP, Nirschl JJ, Klinman E, Holzbaur EL. Amyotrophic lateral sclerosis-linked mutations increase the viscosity of liquid-like TDP-43 RNP granules in neurons. Proc Natl Acad Sci U S A. 2017;114(12):E2466–E75.

17. Conicella AE, Zerze GH, Mittal J, Fawzi NL. ALS Mutations Disrupt Phase Separation Mediated by alpha-Helical Structure in the TDP-43 Low-Complexity C-Terminal Domain. Structure. 2016;24(9):1537–49.

18. Gao J, Wang L, Ren X, Dunn JR, Peters A, Miyagi M, et al. Translational regulation in the brain by TDP-43 phase separation. J Cell Biol. 2021;220(10).

19. Hallegger M, Chakrabarti AM, Lee FCY, Lee BL, Amalietti AG, Odeh HM, et al. TDP-43 condensation properties specify its RNA-binding and regulatory repertoire. Cell. 2021;184(18):4680–96 e22.

20. Grese ZR, Bastos AC, Mamede LD, French RL, Miller TM, Ayala YM. Specific RNA interactions promote TDP-43 multivalent phase separation and maintain liquid properties. EMBO Rep. 2021;22(12):e53632.

21. de Boer EMJ, Orie VK, Williams T, Baker MR, De Oliveira HM, Polvikoski T, et al. TDP-43 proteinopathies: a new wave of neurodegenerative diseases. J Neurol Neurosurg Psychiatry. 2020;92(1):86–95.

22. Arai T, Hasegawa M, Akiyama H, Ikeda K, Nonaka T, Mori H, et al. TDP-43 is a component of ubiquitin-positive tau-negative inclusions in frontotemporal lobar degeneration and amyotrophic lateral sclerosis. Biochem Biophys Res Commun. 2006;351(3):602–11.

23. Neumann M, Sampathu DM, Kwong LK, Truax AC, Micsenyi MC, Chou TT, et al. Ubiquitinated TDP-43 in frontotemporal lobar degeneration and amyotrophic lateral sclerosis. Science. 2006;314(5796):130–3.

24. Amador-Ortiz C, Lin WL, Ahmed Z, Personett D, Davies P, Duara R, et al. TDP-43 immunoreactivity in hippocampal sclerosis and Alzheimer’s disease. Ann Neurol. 2007;61(5):435–45.

25. Nelson PT, Dickson DW, Trojanowski JQ, Jack CR, Boyle PA, Arfanakis K, et al. Limbic-predominant age-related TDP-43 encephalopathy (LATE): consensus working group report. Brain. 2019;142(6):1503–27.

26. McKee AC, Gavett BE, Stern RA, Nowinski CJ, Cantu RC, Kowall NW, et al. TDP-43 proteinopathy and motor neuron disease in chronic traumatic encephalopathy. J Neuropathol Exp Neurol. 2010;69(9):918–29.

27. Brown AL, Wilkins OG, Keuss MJ, Hill SE, Zanovello M, Lee WC, et al. TDP-43 loss and ALS-risk SNPs drive mis-splicing and depletion of UNC13A. Nature. 2022;603(7899):131–7.

28. Estades Ayuso V, Pickles S, Todd T, Yue M, Jansen-West K, Song Y, et al. TDP-43-regulated cryptic RNAs accumulate in Alzheimer’s disease brains. Mol Neurodegener. 2023;18(1):57.

29. Ling JP, Pletnikova O, Troncoso JC, Wong PC. TDP-43 repression of nonconserved cryptic exons is compromised in ALS-FTD. Science. 2015;349(6248):650–5.

30. Ma XR, Prudencio M, Koike Y, Vatsavayai SC, Kim G, Harbinski F, et al. TDP-43 represses cryptic exon inclusion in the FTD-ALS gene UNC13A. Nature. 2022;603(7899):124–30.

31. Melamed Z, Lopez-Erauskin J, Baughn MW, Zhang O, Drenner K, Sun Y, et al. Premature polyadenylation-mediated loss of stathmin-2 is a hallmark of TDP-43-dependent neurodegeneration. Nat Neurosci. 2019;22(2):180–90.

32. Al-Chalabi A, Hardiman O. The epidemiology of ALS: a conspiracy of genes, environment and time. Nat Rev Neurol. 2013;9(11):617–28.

33. Goutman SA, Savelieff MG, Jang DG, Hur J, Feldman EL. The amyotrophic lateral sclerosis exposome: recent advances and future directions. Nat Rev Neurol. 2023;19(10):617–34.

34. Chio A, Mazzini L, D’Alfonso S, Corrado L, Canosa A, Moglia C, et al. The multistep hypothesis of ALS revisited: The role of genetic mutations. Neurology. 2018;91(7):e635–e42.

35. Al-Chalabi A, Calvo A, Chio A, Colville S, Ellis CM, Hardiman O, et al. Analysis of amyotrophic lateral sclerosis as a multistep process: a population-based modelling study. Lancet Neurol. 2014;13(11):1108–13.

36. Barker S, Paul BD, Pieper AA. Increased Risk of Aging-Related Neurodegenerative Disease after Traumatic Brain Injury. Biomedicines. 2023;11(4).

37. Heyburn L, Sajja V, Long JB. The Role of TDP-43 in Military-Relevant TBI and Chronic Neurodegeneration. Front Neurol. 2019;10:680.

38. Jankovic T, Dolenec P, Rajic Bumber J, Grzeta N, Kriz J, Zupan G, et al. Differential Expression Patterns of TDP-43 in Single Moderate versus Repetitive Mild Traumatic Brain Injury in Mice. Int J Mol Sci. 2021;22(22).

39. Kahriman A, Bouley J, Smith TW, Bosco DA, Woerman AL, Henninger N. Mouse closed head traumatic brain injury replicates the histological tau pathology pattern of human disease: characterization of a novel model and systematic review of the literature. Acta Neuropathol Commun. 2021;9(1):118.

40. Kahriman A, Bouley J, Tuncali I, Dogan EO, Pereira M, Luu T, et al. Repeated mild traumatic brain injury triggers pathology in asymptomatic C9ORF72 transgenic mice. Brain. 2023;146(12):5139–52.

41. Wiesner D, Tar L, Linkus B, Chandrasekar A, Olde Heuvel F, Dupuis L, et al. Reversible induction of TDP-43 granules in cortical neurons after traumatic injury. Exp Neurol. 2018;299(Pt A):15–25.

42. Stein TD, Alvarez VE, McKee AC. Chronic traumatic encephalopathy: a spectrum of neuropathological changes following repetitive brain trauma in athletes and military personnel. Alzheimers Res Ther. 2014;6(1):4.

43. Henninger N, Bouley J, Sikoglu EM, An J, Moore CM, King JA, et al. Attenuated traumatic axonal injury and improved functional outcome after traumatic brain injury in mice lacking Sarm1. Brain. 2016;139(Pt 4):1094–105.

44. White MA, Kim E, Duffy A, Adalbert R, Phillips BU, Peters OM, et al. TDP-43 gains function due to perturbed autoregulation in a Tardbp knock-in mouse model of ALS-FTD. Nat Neurosci. 2018;21(4):552–63.

45. Hasan R, Humphrey J, Bettencourt C, Newcombe J, Consortium NA, Lashley T, et al. Transcriptomic analysis of frontotemporal lobar degeneration with TDP-43 pathology reveals cellular alterations across multiple brain regions. Acta Neuropathol. 2022;143(3):383–401.

46. Humphrey J, Venkatesh S, Hasan R, Herb JT, de Paiva Lopes K, Kucukali F, et al. Integrative transcriptomic analysis of the amyotrophic lateral sclerosis spinal cord implicates glial activation and suggests new risk genes. Nat Neurosci. 2023;26(1):150–62.

47. Neelagandan N, Gonnella G, Dang S, Janiesch PC, Miller KK, Kuchler K, et al. TDP-43 enhances translation of specific mRNAs linked to neurodegenerative disease. Nucleic Acids Res. 2019;47(1):341–61.

48. Russo A, Scardigli R, La Regina F, Murray ME, Romano N, Dickson DW, et al. Increased cytoplasmic TDP-43 reduces global protein synthesis by interacting with RACK1 on polyribosomes. Human molecular genetics. 2017;26(8):1407–18.

49. Bjork RT, Mortimore NP, Loganathan S, Zarnescu DC. Dysregulation of Translation in TDP-43 Proteinopathies: Deficits in the RNA Supply Chain and Local Protein Production. Front Neurosci. 2022;16:840357.

50. Piol D, Robberechts T, Da Cruz S. Lost in local translation: TDP-43 and FUS in axonal/neuromuscular junction maintenance and dysregulation in amyotrophic lateral sclerosis. Neuron. 2023;111(9):1355–80.

51. Zhong J, Gunner G, Henninger N, Schafer DP, Bosco DA. Intravital Imaging of Fluorescent Protein Expression in Mice with a Closed-Skull Traumatic Brain Injury and Cranial Window Using a Two-Photon Microscope. J Vis Exp. 2023(194).

52. Dogan EO, Bouley J, Zhong J, Harkins AL, Keeler AM, Bosco DA, et al. Genetic ablation of Sarm1 attenuates expression and mislocalization of phosphorylated TDP-43 after mouse repetitive traumatic brain injury. Acta Neuropathol Commun. 2023;11(1):206.

53. Flierl MA, Stahel PF, Beauchamp KM, Morgan SJ, Smith WR, Shohami E. Mouse closed head injury model induced by a weight-drop device. Nat Protoc. 2009;4(9):1328–37.

54. Kim D, Paggi JM, Park C, Bennett C, Salzberg SL. Graph-based genome alignment and genotyping with HISAT2 and HISAT-genotype. Nat Biotechnol. 2019;37(8):907–15.

55. Chen Y, Lun AT, Smyth GK. From reads to genes to pathways: differential expression analysis of RNA-Seq experiments using Rsubread and the edgeR quasi-likelihood pipeline. F1000Res. 2016;5:1438.

56. Castanza AS, Recla JM, Eby D, Thorvaldsdottir H, Bult CJ, Mesirov JP. Extending support for mouse data in the Molecular Signatures Database (MSigDB). Nat Methods. 2023;20(11):1619–20.

57. Liberzon A, Birger C, Thorvaldsdottir H, Ghandi M, Mesirov JP, Tamayo P. The Molecular Signatures Database (MSigDB) hallmark gene set collection. Cell Syst. 2015;1(6):417–25.

58. Liberzon A, Subramanian A, Pinchback R, Thorvaldsdottir H, Tamayo P, Mesirov JP. Molecular signatures database (MSigDB) 3.0. Bioinformatics. 2011;27(12):1739–40.

59. Mootha VK, Lindgren CM, Eriksson KF, Subramanian A, Sihag S, Lehar J, et al. PGC-1alpha-responsive genes involved in oxidative phosphorylation are coordinately downregulated in human diabetes. Nat Genet. 2003;34(3):267–73.

60. Subramanian A, Tamayo P, Mootha VK, Mukherjee S, Ebert BL, Gillette MA, et al. Gene set enrichment analysis: a knowledge-based approach for interpreting genome-wide expression profiles. Proc Natl Acad Sci U S A. 2005;102(43):15545–50.

61. Funes S, Jung J, Gadd DH, Mosqueda M, Zhong J, Shankaracharya, et al. Expression of ALS-PFN1 impairs vesicular degradation in iPSC-derived microglia. Nat Commun. 2024;15(1):2497.

62. Tyanova S, Temu T, Cox J. The MaxQuant computational platform for mass spectrometry-based shotgun proteomics. Nat Protoc. 2016;11(12):2301–19.

63. Tyanova S, Temu T, Sinitcyn P, Carlson A, Hein MY, Geiger T, et al. The Perseus computational platform for comprehensive analysis of (prote)omics data. Nat Methods. 2016;13(9):731–40.

64. Schindelin J, Arganda-Carreras I, Frise E, Kaynig V, Longair M, Pietzsch T, et al. Fiji: an open-source platform for biological-image analysis. Nat Methods. 2012;9(7):676–82.

65. Kamentsky L, Jones TR, Fraser A, Bray MA, Logan DJ, Madden KL, et al. Improved structure, function and compatibility for CellProfiler: modular high-throughput image analysis software. Bioinformatics. 2011;27(8):1179–80.

66. Jumper J, Evans R, Pritzel A, Green T, Figurnov M, Ronneberger O, et al. Highly accurate protein structure prediction with AlphaFold. Nature. 2021;596(7873):583-9.

67. Darnell JC, Van Driesche SJ, Zhang C, Hung KY, Mele A, Fraser CE, et al. FMRP stalls ribosomal translocation on mRNAs linked to synaptic function and autism. Cell. 2011;146(2):247–61.

68. Dogan EO, Simonini SR, Bouley J, Weiss A, Brown RH, Jr., Henninger N. Genetic Ablation of Sarm1 Mitigates Disease Acceleration after Traumatic Brain Injury in the SOD1(G93A) Transgenic Mouse Model of Amyotrophic Lateral Sclerosis. Ann Neurol. 2025.

69. Curzon P, Rustay NR, Browman KE. Cued and Contextual Fear Conditioning for Rodents. 2nd ed: CRC Press/Taylor & Francis, Boca Raton (FL); 2009 2009.

70. Maren S, Quirk GJ. Neuronal signalling of fear memory. Nat Rev Neurosci. 2004;5(11):844–52.

71. Querfurth H, Lee HK. Mammalian/mechanistic target of rapamycin (mTOR) complexes in neurodegeneration. Mol Neurodegener. 2021;16(1):44.

72. Simon DW, McGeachy MJ, Bayir H, Clark RS, Loane DJ, Kochanek PM. The far-reaching scope of neuroinflammation after traumatic brain injury. Nat Rev Neurol. 2017;13(3):171–91.

73. Hubbard WB, Joseph B, Spry M, Vekaria HJ, Saatman KE, Sullivan PG. Acute Mitochondrial Impairment Underlies Prolonged Cellular Dysfunction after Repeated Mild Traumatic Brain Injuries. J Neurotrauma. 2019;36(8):1252–63.

74. Wang W, Wang L, Lu J, Siedlak SL, Fujioka H, Liang J, et al. The inhibition of TDP-43 mitochondrial localization blocks its neuronal toxicity. Nat Med. 2016;22(8):869–78.

75. Yoshida H, Matsui T, Yamamoto A, Okada T, Mori K. XBP1 mRNA is induced by ATF6 and spliced by IRE1 in response to ER stress to produce a highly active transcription factor. Cell. 2001;107(7):881–91.

76. Pineda SS, Lee H, Ulloa-Navas MJ, Linville RM, Garcia FJ, Galani K, et al. Single-cell dissection of the human motor and prefrontal cortices in ALS and FTLD. Cell. 2024;187(8):1971–89 e16.

77. Perucho L, Artero-Castro A, Guerrero S, Ramon y Cajal S, Me LL, Wang ZQ. RPLP1, a crucial ribosomal protein for embryonic development of the nervous system. PLoS One. 2014;9(6):e99956.

78. Moisse K, Volkening K, Leystra-Lantz C, Welch I, Hill T, Strong MJ. Divergent patterns of cytosolic TDP-43 and neuronal progranulin expression following axotomy: implications for TDP-43 in the physiological response to neuronal injury. Brain Res. 2009;1249:202–11.

79. Mehta PR, Brown AL, Ward ME, Fratta P. The era of cryptic exons: implications for ALS-FTD. Mol Neurodegener. 2023;18(1):16.

80. Rouaux C, Gonzalez De Aguilar JL, Dupuis L. Unmasking the skiptic task of TDP-43. EMBO J. 2018;37(11).

81. Arnold ES, Ling SC, Huelga SC, Lagier-Tourenne C, Polymenidou M, Ditsworth D, et al. ALS-linked TDP-43 mutations produce aberrant RNA splicing and adult-onset motor neuron disease without aggregation or loss of nuclear TDP-43. Proc Natl Acad Sci U S A. 2013;110(8):E736–45.

82. Fratta P, Sivakumar P, Humphrey J, Lo K, Ricketts T, Oliveira H, et al. Mice with endogenous TDP-43 mutations exhibit gain of splicing function and characteristics of amyotrophic lateral sclerosis. EMBO J. 2018;37(11).

83. Polymenidou M, Lagier-Tourenne C, Hutt KR, Huelga SC, Moran J, Liang TY, et al. Long pre-mRNA depletion and RNA missplicing contribute to neuronal vulnerability from loss of TDP-43. Nat Neurosci. 2011;14(4):459–68.

84. Yang C, Qiao T, Yu J, Wang H, Guo Y, Salameh J, et al. Low-level overexpression of wild type TDP-43 causes late-onset, progressive neurodegeneration and paralysis in mice. PLoS One. 2022;17(2):e0255710.

85. Chu BW, Kovary KM, Guillaume J, Chen LC, Teruel MN, Wandless TJ. The E3 ubiquitin ligase UBE3C enhances proteasome processivity by ubiquitinating partially proteolyzed substrates. J Biol Chem. 2013;288(48):34575–87.

86. Gaetani M, Zubarev RA. Proteome Integral Solubility Alteration (PISA) for High-Throughput Ligand Target Deconvolution with Increased Statistical Significance and Reduced Sample Amount. Methods Mol Biol. 2023;2554:91–106.

87. Li J, Van Vranken JG, Paulo JA, Huttlin EL, Gygi SP. Selection of Heating Temperatures Improves the Sensitivity of the Proteome Integral Solubility Alteration Assay. J Proteome Res. 2020;19(5):2159–66.

88. Savitski MM, Reinhard FB, Franken H, Werner T, Savitski MF, Eberhard D, et al. Tracking cancer drugs in living cells by thermal profiling of the proteome. Science. 2014;346(6205):1255784.

89. Brohee S, van Helden J. Evaluation of clustering algorithms for protein-protein interaction networks. BMC Bioinformatics. 2006;7:488.

90. Szklarczyk D, Kirsch R, Koutrouli M, Nastou K, Mehryary F, Hachilif R, et al. The STRING database in 2023: protein-protein association networks and functional enrichment analyses for any sequenced genome of interest. Nucleic Acids Res. 2023;51(D1):D638–D46.

91. Cheng F, Chapman T, Venturato J, Davidson JM, Polido SA, Rosa-Fernandes L, et al. Proteomics Analysis of the TDP-43 Interactome in Cellular Models of ALS Pathogenesis. J Neurochem. 2025;169(5):e70079.

92. Freibaum BD, Chitta RK, High AA, Taylor JP. Global analysis of TDP-43 interacting proteins reveals strong association with RNA splicing and translation machinery. J Proteome Res. 2010;9(2):1104–20.

93. Zeng C, Han S, Pan Y, Huang Z, Zhang B, Zhang B. Revisiting the chaperonin T-complex protein-1 ring complex in human health and disease: A proteostasis modulator and beyond. Clin Transl Med. 2024;14(2):e1592.

94. Luo J, Zhao H, Chen L, Liu M. Multifaceted functions of RPS27a: An unconventional ribosomal protein. J Cell Physiol. 2023;238(3):485–97.

95. Lin Z, Kim E, Ahmed M, Han G, Simmons C, Redhead Y, et al. MRI-guided histology of TDP-43 knock-in mice implicates parvalbumin interneuron loss, impaired neurogenesis and aberrant neurodevelopment in amyotrophic lateral sclerosis-frontotemporal dementia. Brain Commun. 2021;3(2):fcab114.

96. Kelmer Sacramento E, Kirkpatrick JM, Mazzetto M, Baumgart M, Bartolome A, Di Sanzo S, et al. Reduced proteasome activity in the aging brain results in ribosome stoichiometry loss and aggregation. Mol Syst Biol. 2020;16(6):e9596.

97. Amirbeigiarab S, Kiani P, Velazquez Sanchez A, Krisp C, Kazantsev A, Fester L, et al. Invariable stoichiometry of ribosomal proteins in mouse brain tissues with aging. Proc Natl Acad Sci U S A. 2019;116(45):22567–72.

98. Zhao B, Cowan CM, Coutts JA, Christy DD, Saraph A, Hsueh SCC, et al. Targeting RACK1 to alleviate TDP-43 and FUS proteinopathy-mediated suppression of protein translation and neurodegeneration. Acta Neuropathol Commun. 2023;11(1):200.

99. Bhavsar RB, Makley LN, Tsonis PA. The other lives of ribosomal proteins. Hum Genomics. 2010;4(5):327–44.

100. Simsek D, Barna M. An emerging role for the ribosome as a nexus for post-translational modifications. Curr Opin Cell Biol. 2017;45:92–101.

101. Ali A, Garde R, Schaffer OC, Bard JAM, Husain K, Kik SK, et al. Adaptive preservation of orphan ribosomal proteins in chaperone-dispersed condensates. Nat Cell Biol. 2023;25(11):1691–703.

102. Nagano S, Jinno J, Abdelhamid RF, Jin Y, Shibata M, Watanabe S, et al. TDP-43 transports ribosomal protein mRNA to regulate axonal local translation in neuronal axons. Acta Neuropathol. 2020;140(5):695–713.

103. Tank EM, Figueroa-Romero C, Hinder LM, Bedi K, Archbold HC, Li X, et al. Abnormal RNA stability in amyotrophic lateral sclerosis. Nat Commun. 2018;9(1):2845.

104. Chou A, Krukowski K, Jopson T, Zhu PJ, Costa-Mattioli M, Walter P, et al. Inhibition of the integrated stress response reverses cognitive deficits after traumatic brain injury. Proc Natl Acad Sci U S A. 2017;114(31):E6420–E6.

105. Huang WP, Ellis BCS, Hodgson RE, Sanchez Avila A, Kumar V, Rayment J, et al. Stress-induced TDP-43 nuclear condensation causes splicing loss of function and STMN2 depletion. Cell Rep. 2024;43(7):114421.

106. Rifai OM, Waldron FM, O’Shaughnessy J, Read FL, Gilodi M, Pastore A, et al. Amygdala TDP-43 pathology is associated with behavioural dysfunction and ferritin accumulation in amyotrophic lateral sclerosis. bioRxiv. 2024.

107. Wilkins OG, Chien M, Wlaschin JJ, Barattucci S, Harley P, Mattedi F, et al. Creation of de novo cryptic splicing for ALS and FTD precision medicine. Science. 2024;386(6717):61–9.

108. Francois-Moutal L, Perez-Miller S, Scott DD, Miranda VG, Mollasalehi N, Khanna M. Structural Insights Into TDP-43 and Effects of Post-translational Modifications. Front Mol Neurosci. 2019;12:301.

109. Shenouda M, Xiao S, MacNair L, Lau A, Robertson J. A C-Terminally Truncated TDP-43 Splice Isoform Exhibits Neuronal Specific Cytoplasmic Aggregation and Contributes to TDP-43 Pathology in ALS. Front Neurosci. 2022;16:868556.

110. Elliott E, Bailey O, Waldron FM, Hardingham GE, Chandran S, Gregory JM. Therapeutic Targeting of Proteostasis in Amyotrophic Lateral Sclerosis-a Systematic Review and Meta-Analysis of Preclinical Research. Front Neurosci. 2020;14:511.

